# BHLHE40/41 regulate macrophage/microglia responses associated with Alzheimer’s disease and other disorders of lipid-rich tissues

**DOI:** 10.1101/2023.02.13.528372

**Authors:** Anna Podlesny-Drabiniok, Gloriia Novikova, Yiyuan Liu, Josefine Dunst, Rose Temizer, Chiara Giannarelli, Samuele Marro, Taras Kreslavsky, Edoardo Marcora, Alison Mary Goate

**Affiliations:** Department of Genetics and Genomic Sciences, New York, NY, USA; Icahn School of Medicine at Mount Sinai, New York, NY, USA; OMNI Bioinformatics Department and Neuroscience Department, _Genentech_, _Inc._, _South_ San Francisco, CA, USA; Department of Medicine, Division of Immunology and Allergy, Karolinska Institutet, Karolinska University Hospital, Stockholm, Sweden; Center for Molecular Medicine, Karolinska Institutet, Stockholm, Sweden; Department of Medicine (C.G.), Cardiology, NYU Grossman School of Medicine.; Department of Pathology (C.G.), Cardiology, NYU Grossman School of Medicine.; Nash Family Department of Neuroscience, Friedman Brain Institute, Icahn School of Medicine at Mount Sinai, New York, NY 10029, USA; Black Family Stem Cell Institute, Icahn School of Medicine at Mount Sinai, New York, NY 10029, USA

## Abstract

**Background:** Genetic and experimental evidence strongly implicates myeloid cells in the etiology of AD and suggests that AD-associated alleles and genes may modulate disease risk by altering the transcriptional and cellular responses of macrophages (like microglia) to damage of lipid-rich tissues (like the brain). Specifically, recent single-cell/nucleus RNA sequencing (sc/nRNA-seq) studies identified a transcriptionally distinct state of subsets of macrophages in aging or degenerating brains (usually referred to as disease-associated microglia or DAM) and in other diseased lipid-rich tissues (e.g., obese adipose tissue, fatty liver, and atherosclerotic plaques). We collectively refer to these subpopulations as lipid-associated macrophages or LAMs. Importantly, this particular activation state is characterized by increased expression of genes involved in the phagocytic clearance of lipid-rich cellular debris (efferocytosis), including several AD risk genes.

**Methods:** We used sc/nRNA-seq data from human and mouse microglia from healthy and diseased brains and macrophages from other lipid-rich tissues to reconstruct gene regulatory networks and identify transcriptional regulators whose regulons are enriched for LAM response genes (LAM TFs) across species. We then used gene knock- down/knock-out strategies to validate some of these LAM TFs in human THP-1 macrophages and iPSC-derived microglia *in vitro*, as well as mouse microglia *in vivo*.

**Results:** We nominate 11 strong candidate LAM TFs shared across human and mouse networks (*BHLHE41*, *HIF1A*, *ID2*, *JUNB*, *MAF*, *MAFB*, *MEF2A*, *MEF2C*, *NACA, POU2F2* and *SPI1*). We also demonstrate a strong enrichment of AD risk alleles in the cistrome of *BHLHE41* (and its close homolog *BHLHE40*), thus implicating its regulon in the modulation of disease susceptibility. Loss or reduction of *BHLHE40/41* expression in human THP-1 macrophages and iPSC-derived microglia, as well as loss of *Bhlhe40*/*41* in mouse microglia led to increased expression of LAM response genes, specifically those involved in cholesterol clearance and lysosomal processing, with a concomitant increase in cholesterol efflux and storage, as well as lysosomal mass and degradative capacity.

**Conclusions:** Taken together, this study nominates transcriptional regulators of the LAM response, experimentally validates BHLHE40/41 in human and mouse macrophages/microglia, and provides novel targets for therapeutic modulation of macrophage/microglia function in AD and other disorders of lipid-rich tissues.

**Graphical abstract:** 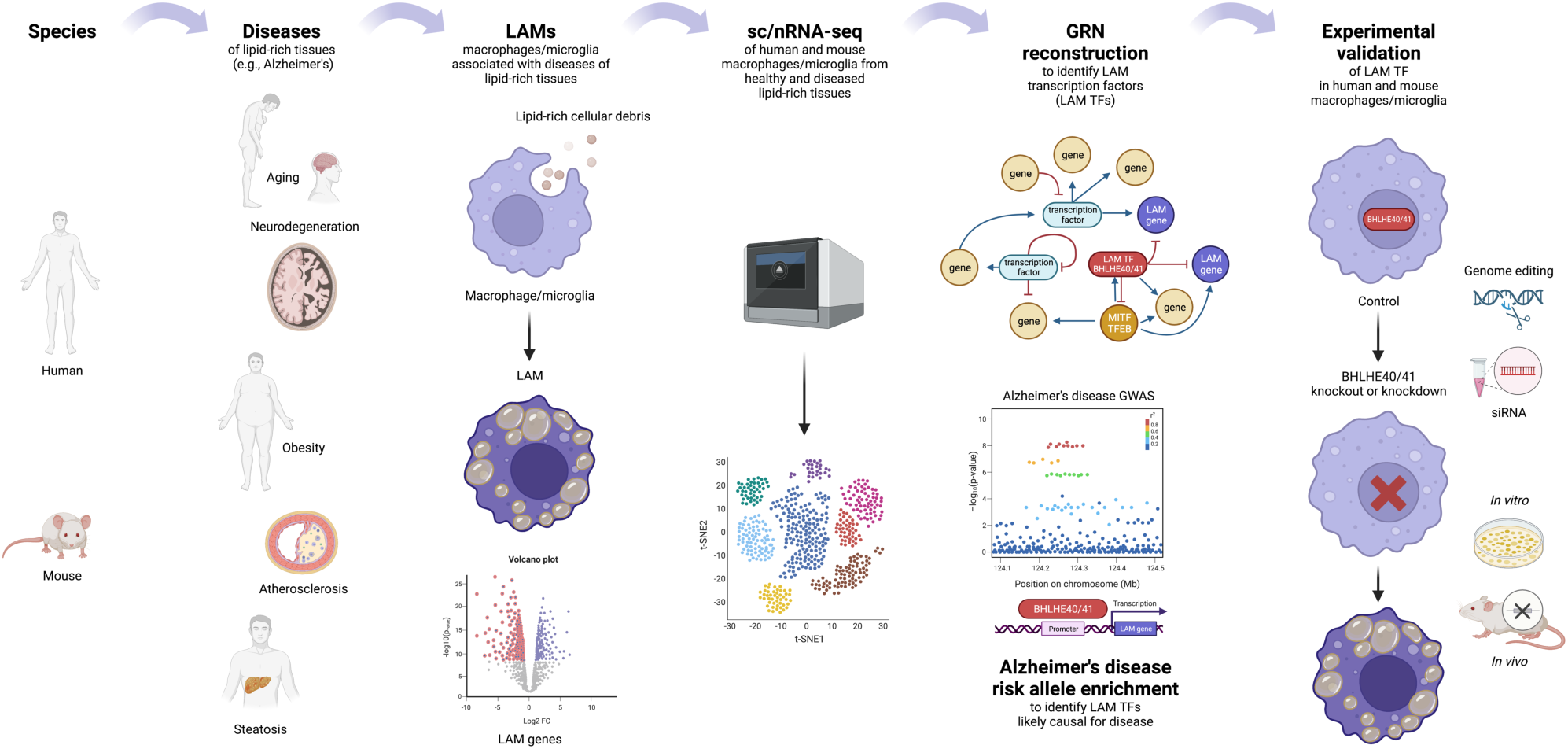

## Introduction

Tissue-resident and monocyte-derived macrophages are myeloid cells that specialize in the phagocytic clearance of host tissue-derived cellular debris (efferocytosis). This core function of macrophages is essential for the maintenance of tissue homeostasis and immune tolerance, the resolution of inflammation, and the repair of damaged tissue. Macrophage dysfunction has been implicated in aging and the pathogenesis of numerous diseases, including Alzheimer’s disease (AD), demyelinating disorders, schizophrenia, obesity, steatosis, atherosclerosis, and several other autoimmune and chronic inflammatory diseases [1–6].

In AD, genetic and experimental evidence strongly implicates myeloid cells (including microglia, the brain-resident macrophages) in disease pathogenesis and suggests that AD-associated alleles and genes may modulate disease risk by altering the transcriptional and cellular responses of macrophages/microglia to damaged cholesterol/lipid-rich tissues (such as the aging or degenerating brain, the most cholesterol/lipid-rich organ of the human body [7]). Specifically, recent single-cell/nucleus RNA sequencing (sc/nRNA-seq) studies identified a transcriptionally distinct state of subsets of macrophages in aging or degenerating brains (usually referred to as disease-associated microglia or DAM) and in other diseased lipid-rich tissues (e.g., obese adipose tissue, fatty liver, and atherosclerotic plaques). In this manuscript, we refer to these subpopulations as lipid-associated macrophages or LAMs.

Importantly, this particular activation state is characterized by increased expression of efferocytosis (e.g., phagocytic clearance and cholesterol/lipid metabolism) genes, including several AD risk genes [8, 9]. Some of these AD risk genes (*TREM2*, *PLCG2*, and *APOE*) are critical for the development and function of LAMs in response to cholesterol/lipid overload in mouse models of AD, demyelination, and other disorders of lipid-rich tissues [10–13]. Therefore, a deeper understanding of the transcriptional regulation of the LAM response may offer valuable biological insights and aid the prioritization of novel therapeutic targets for AD and other disorders of lipid-rich tissues.

The LAM response, like other transcriptional and cellular responses, is likely regulated by transcription factors (TFs) that orchestrate the coordinated expression of LAM effector genes through direct binding to their promoters and other *cis*-regulatory elements. Reconstruction of TF-centric gene regulatory networks (GRNs) can lead to the identification of TF target genes and co-regulated gene modules (regulons) [14]. In turn, enrichment of LAM signature genes in these regulons can point to candidate TFs that are likely to direct this response [14]. Integration of multiple microglial bulk microarray datasets, including transcriptome profiles of microglial cells acutely isolated from AD mouse brains or cultured and treated with LPS or IL4, led to the identification of potential regulators of several microglial transcriptional states and identified *Cebpα*, *Irf1*, and *Lxrα/β* as positive transcriptional regulators of LAM markers such as *Apoe, Cxcr4,* and *Trem2* [15, 16]. Another group characterized microglia co-expression modules and found that *Bhlhe40, Rxrγ, Hif1*α and *Mitf* were coexpressed with genes enriched in modules of neurodegeneration-related microglia that resemble the LAM transcriptional state [17].

Here, we used sc/nRNA-seq data of human and mouse macrophages from healthy and diseased brains and other lipid-rich tissues to reconstruct GRNs and identify TFs whose regulons are enriched for LAM signature genes (LAM TFs) across species. We nominate 11 strong candidate LAM TFs shared across human and mouse networks (*BHLHE41*, *HIF1A*, *ID2*, *JUNB*, *MAF*, *MAFB*, *MEF2A*, *MEF2C*, *NACA, POU2F2* and *SPI1*). We also demonstrate a strong enrichment of AD risk alleles in the cistrome of *BHLHE41* (and its close homolog *BHLHE40*), thus implicating its regulon in the modulation of disease susceptibility. Loss or reduction of *BHLHE40/41* expression in human THP-1 macrophages and iPSC-derived microglia as well as loss of *Bhlhe40*/*41* in mouse microglia led to increased expression of a subset of LAM response genes, specifically those involved in cholesterol clearance and lysosomal processing, with a concomitant increase in cholesterol efflux and storage, as well as lysosomal mass and degradative capacity. Taken together, this study nominates transcriptional regulators of the LAM response, experimentally validates BHLHE40/41 in human and mouse macrophages, and provides novel targets for therapeutic modulation of macrophage/microglia function in AD and other disorders of lipid-rich tissues.

## Results

### Reconstruction of gene regulatory networks in human and mouse macrophages/microglia using single-cell/nucleus RNA sequencing data

To reconstruct GRNs and nominate candidate transcriptional regulators of the LAM response (henceforth referred to as LAM TFs), we used eight human sc/nRNA-seq and ten mouse scRNA-seq macrophage/microglia datasets from healthy and diseased brains and other lipid-rich tissues (Methods, Supplementary Table 1). Human datasets included microglia from healthy and AD brains, macrophages from atherosclerotic plaques, macrophages from adipose tissue of healthy and obese individuals, and Kupffer cells from healthy and cirrhotic livers [3,18–24]. Mouse datasets included microglia from control mice and mouse models of amyloid deposition (i.e AppNL-G-F, APP/PS1 and 5XFAD), microglia from aging and demyelinating brains (i.e., upon cuprizone treatment), macrophages from atherosclerotic plaques, macrophages from adipose tissue of healthy and obese mice, and Kupffer cells from healthy, non-alcoholic steatohepatitis (NASH), and cirrhotic livers [3,12,18,19,25–29]. GRNs were reconstructed by computing mutual information between annotated transcriptional regulators (642 human and 641 mouse) and all other expressed genes in each dataset using ARACNe [30] (Methods). To filter out likely indirect TF-target interactions, we used data processing inequality (DPI) in ARACNe, thus enriching the regulon for potential direct targets [30] (Methods).

### Geneset enrichment analysis of TF regulons nominates candidate LAM TFs in human and mouse macrophages/microglia

To nominate candidate LAM TFs, we tested the regulons of all TFs in human and mouse GRNs for over-representation of LAM signature genes reported in 1) DAM microglia from brains of 5XFAD mice (Table S2 from Keren-Shaul *et al*. [10]); 2) TREM2^hi^ macrophages from atherosclerotic plaques of *Ldlr^-/-^* mice on high-fat diet (Online Table I, cluster 11 from Cochain *et al*. [26]) and 3) LAM macrophages from human and mouse obese adipose tissue (Dataset S6 and Dataset S5 from Jaitin *et al*. [3]) (Methods and Supplementary Table 1). This approach allowed us to prioritize TFs whose regulons displayed a statistically significant enrichment of LAM signature genes, suggesting they may at least partially regulate these transcriptional responses. Since LAM response genes are partially conserved in various macrophage types and across species [24], we focused on TFs that were nominated in both human and mouse networks and whose regulons were enriched for all three LAM genesets mentioned above. Seventy-four LAM TFs were nominated in at least two mouse and two human GRNs using all three LAM genesets (Figure 1A-B).

**Figure 1.**
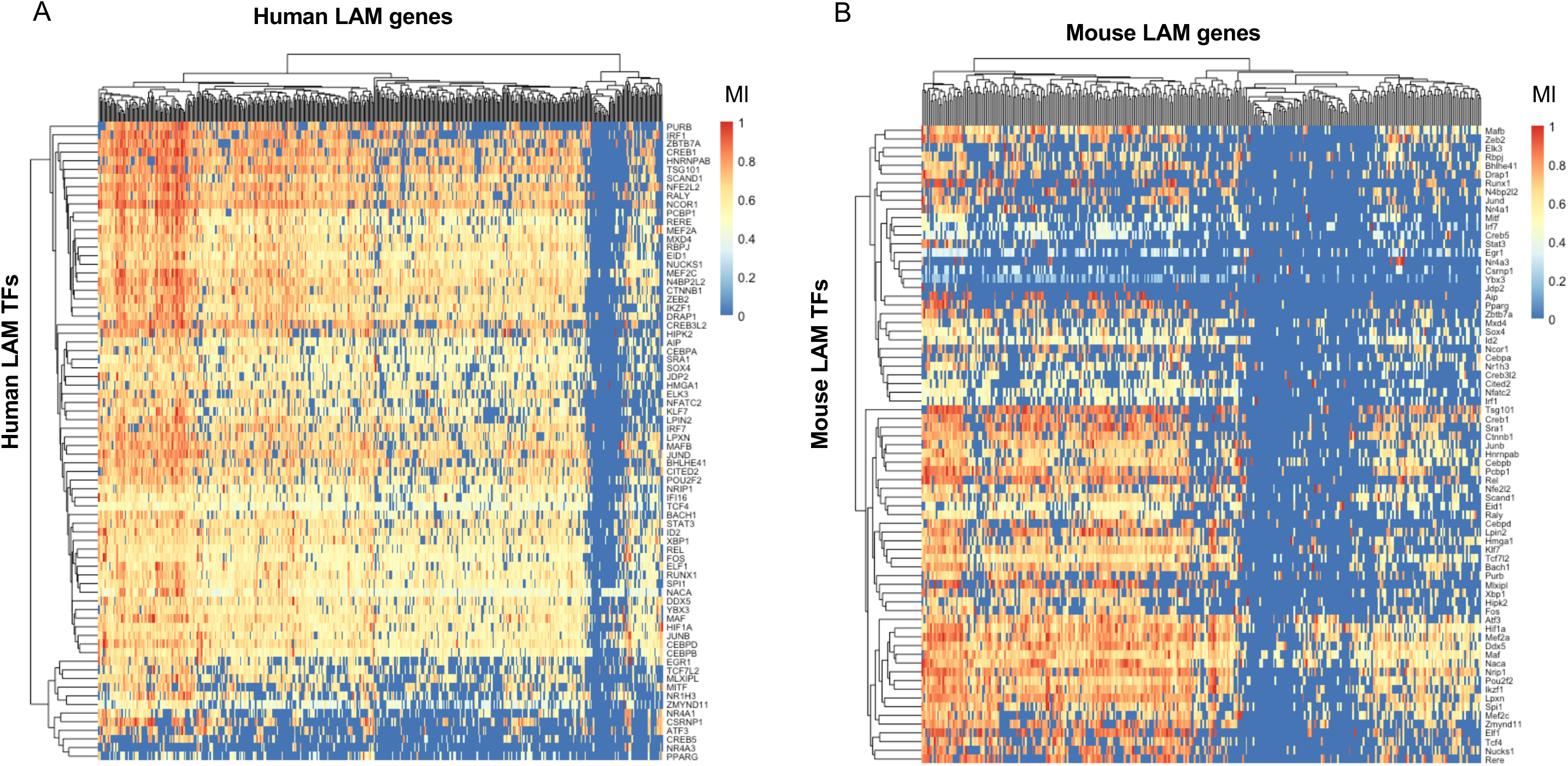
Gene regulatory network analysis of sc/snRNA-Seq datasets from human and mouse macrophages/microglia nominates 74 transcription factors as candidate transcriptional regulators of the lipid-associated macrophage (LAM) response (LAM TFs). **A)** Heatmap showing normalized mutual interaction (MI) values for each LAM gene - LAM transcription factor (LAM TF) pair in a meta- analyzed human network (8 human networks were meta-analyzed). Human LAM genes are from Jaitin *et al*. [3] (Dataset S6, FDR Adj.P- value < 0.05). **B)** Heatmap showing normalized MI values for each LAM gene - LAM TF pair in a meta-analyzed mouse network (10 mouse networks were meta-analyzed). Mouse LAM genes are from Keren-Shaul *et al*. [10] (Table S2). The 74 LAM TFs were nominated in at least two human and mouse networks, were conserved between species, and their regulons were significantly enriched for all three LAM genesets (DAM, LAM and TREM2^hi^, Supplementary Table 1).

To generate a shortlist of LAM TFs, we required each transcription factor to be nominated in at least half of human and mouse networks and to be expressed in human microglia [31]. Using this more stringent set of criteria, we nominated 11 genes (*BHLHE41*, *HIF1A*, *ID2*, *JUNB*, *MAF*, *MAFB*, *MEF2A*, *MEF2C*, *NACA, POU2F2* and *SPI1*) as strong candidate LAM TFs in both human and mouse networks for further analyses. We were unable to identify a candidate transcriptional regulator for a small number of LAM response genes, while some LAM response genes were predicted to be regulated by only a handful of TFs in both human and mouse networks (Figure 1A-B). A potential explanation for this observation could be that the low expression of these genes in macrophages/microglia prevented us from identifying robust associations. Indeed, some of these genes, such as *CD5*, *CXCL14* and *KIF1A*, have very low expression in microglia [31] despite being significantly upregulated in DAM microglia from 5XFAD mice [10, 31].

Although we focused on TFs nominated in both human and mouse networks for our downstream analyses, we also identified TFs specifically nominated as LAM transcriptional regulators in one species but not the other (i.e., nominated in half of the mouse networks, but not in human networks and vice versa). For example, *Bhlhe40, Atf3,* and *Fli1* are nominated in mouse networks, while *CEBPD, CEBPB*, and *EGR1* are nominated in human networks (Supplementary Table 1). This is in agreement with a pre- print study that recently nominated *CEBPB, CEBPD*, and *EGR1* as candidate drivers of a LAM-like transcriptional state observed in cultured human iPSC-derived microglia (iMGLs) exposed to cholesterol/lipid-rich cellular debris *in vitro* [32]. These differences between species could be driven by true differences in LAM gene expression regulation, differences in the expression levels of these TFs, as well as technical differences between human and mouse datasets. Nevertheless, taken together, these analyses identified conserved and species-specific putative transcriptional regulators of the LAM response.

### Human genetics implicates candidate LAM TFs BHLHE40/41 and their cistromes in the etiology of AD

Several AD risk genes (*APOE*, *PLCG2* and *TREM2*) have been shown to be critical for the development and function of LAMs in response to lipid overload in mouse models of AD and demyelination, and other diseases of lipid-rich tissues, highlighting a potential causal link between regulation of the LAM response and modulation of AD risk [10–13]. Therefore, we investigated whether any of the candidate transcriptional regulators of the LAM response (LAM TFs) nominated in this study could also be implicated in the modulation of AD risk. We previously showed that AD risk alleles are significantly enriched in the cistrome of PU.1 (encoded by *SPI1*, a candidate AD risk gene [33] as well as one of the candidate LAM TFs nominated in this study), suggesting a role for SPI1/PU.1 regulated genes in the etiology of AD [33]. Since then, this finding has been independently replicated multiple times [34, 35]. To investigate whether any of the other candidate LAM TFs exhibit a similar enrichment, we used stratified LD score regression to perform partitioned AD heritability analysis using open chromatin regions (ATAC-seq peaks) that contain binding motifs for the TF of interest (henceforth referred to as TF proxy-binding sites) in microglia and macrophages, as previously described [36]. We could not perform these analyses for *NACA* and *ID2*, since these transcriptional regulators do not have a DNA binding motif in the HOMER database or lack a DNA binding domain [37]. We replicated the significant enrichment of AD risk alleles in SPI1/PU.1 proxy-binding sites in human microglia (P-value = 6.9e-03) and monocyte-derived macrophages (P-value = 1.9e-03) (Figure 2A and B). Our approach using TF proxy-binding sites for partitioned heritability analysis has been previously validated by estimating a similar enrichment of AD risk alleles in SPI1/PU.1 binding sites obtained from SPI1/PU.1 ChIP-seq (as opposed to ATAC-seq) peaks in human monocytes and macrophages [33].

**Figure 2.**
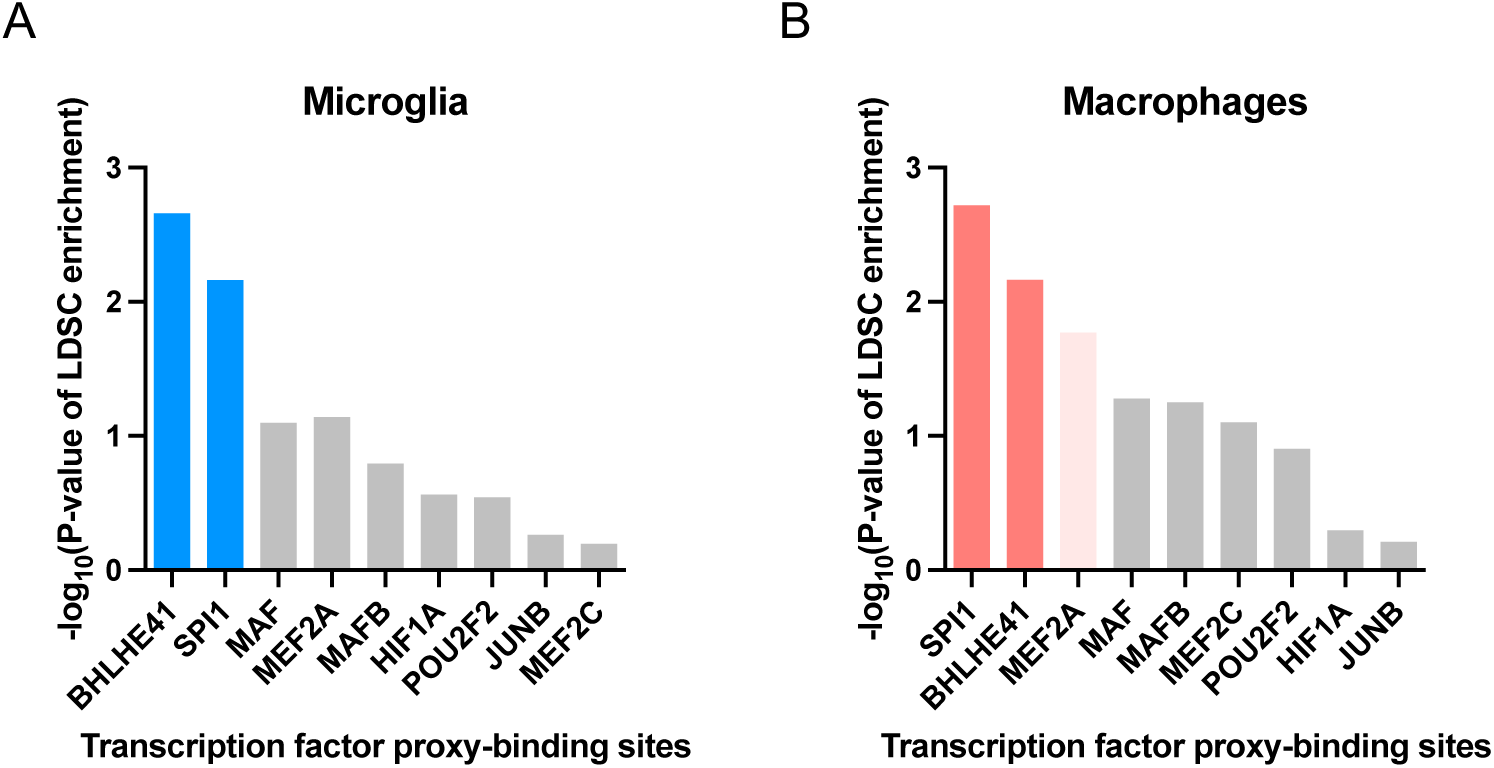
Alzheimer’s disease (AD) risk alleles are enriched in the BHLHE41 and SPI1/PU.1 cistromes. **A)** -log10 of enrichment P-values obtained from stratified LD Score Regression (LDSC) analysis of AD GWAS SNP heritability partitioned by transcription factor (TF) proxy-binding sites, which were obtained by stratifying ATAC-Seq peaks in human microglia by the presence of binding motifs for each candidate LAM TF listed on the x-axis. **B)** -log10 of enrichment P-values obtained from stratified LDSC analysis of AD GWAS SNP heritability partitioned by TF proxy-binding sites, which were obtained by stratifying ATAC-Seq peaks in human monocyte-derived macrophages by the presence of binding motifs for each candidate LAM TF listed on the x-axis. Bars in dark blue or dark red indicate significant enrichments (FDR Adj.P-value < 0.05), bars in light red indicate nominally significant enrichments (P-value < 0.05), while gray bars indicate non-significant enrichments.

In addition to SPI1/PU.1, we uncovered a significant enrichment of AD risk alleles in the proxy-cistrome of BHLHE41 in human microglia and monocyte-derived macrophages (P-value = 2.2e-03 and 6.8e-03, respectively), implicating its target gene network in AD risk modification (Figures 2A and B). In monocyte-derived macrophages, AD risk alleles were also nominally enriched in the proxy-cistrome of MEF2A (P-value = 0.02) (Figure 2B).

Interestingly, a recent AD GWAS in African-Americans uncovered a genome-wide significant locus near *BHLHE40*, a close homolog of *BHLHE41* [38]. Due to the high DNA binding motif similarity between BHLHE40 and BHLHE41 (similarity score of 0.99 in HOMER motif database), BHLHE40 proxy-binding sites were also enriched in AD heritability in both microglia (P-value = 3.7e-03) and monocyte-derived macrophages (P-value = 5.8e-03). BHLHE40 and BHLHE41 (also known as DEC1/SHARP2/BHLHB2/STRA13 and DEC2/SHARP1/BHLHB3, respectively) belong to a family of basic helix-loop-helix transcriptional regulators that counteract transcriptional responses induced by other transcription factors like LXR:RXR nuclear receptors and members of the MiT/TFE family, which includes transcription factor EB (TFEB) and microphthalmia-associated transcription factor (MITF) [39–41]. In addition, BHLHE40 and BHLHE41 often exhibit functional redundancy [42–44]. Taken together, these results suggest that BHLHE40/41 and their target gene networks may modulate AD risk through their ability to regulate the LAM response in macrophages/microglia.

### BHLHE41 and SPI1/PU.1 likely regulate LAM response genes through direct binding to their promoters

Regulation of gene expression usually involves an interplay between multiple proximal and distal cis-regulatory elements and the gene promoter, mediated by specific three-dimensional chromatin arrangements [45]. These gene regulatory elements contain short and specific DNA sequences (motifs) to which TFs can bind to promote or repress transcription [45]. To further refine our list of candidate LAM TFs, we investigated which of them likely regulate their predicted target genes directly by binding to their promoters. To address this question we used open chromatin (ATAC-seq) profiles of human and mouse microglia (Methods). First, for each LAM TF, we quantified the number of instances of the corresponding binding motifs in open chromatin regions (ATAC-seq peaks) within the promoters of genes belonging to the LAM TF regulon. We observed prominent peaks in *BHLHE41* and *SPI1* motif instances, suggesting our GRN reconstruction approach was indeed able to identify regulons that are enriched for direct targets of these LAM TFs (Supplementary Figure 1A and B).

To further explore the question of LAM gene expression regulation, we stratified ATAC-seq peaks in human and mouse microglia by the presence of LAM TF binding motifs and quantified the proportion of LAM signature genes (Jaitin et al. [3], Dataset S6, FDR Adj.P-value < 0.05 for human LAM signature genes; Keren-Shaul *et al.* [10], Table S3, FDR Adj.P-value < 0.05 for mouse LAM signature genes) that contain LAM TF proxy- binding sites in their promoters. BHLHE41 and SPI1/PU.1 proxy-binding sites were observed in the highest proportion of LAM gene promoters in both human and mouse microglia and monocyte-derived macrophages (Figure 3A–D, Supplementary Figure 1A and B). This suggests that BHLHE41 and SPI1/PU.1 likely act as direct regulators of the LAM transcriptional response, while other TFs identified by our analyses with a smaller proportion of proxy-binding sites in LAM promoters might regulate fewer LAM genes and/or regulate them indirectly or by binding to other gene regulatory elements (e.g., enhancers).

**Figure 3.**
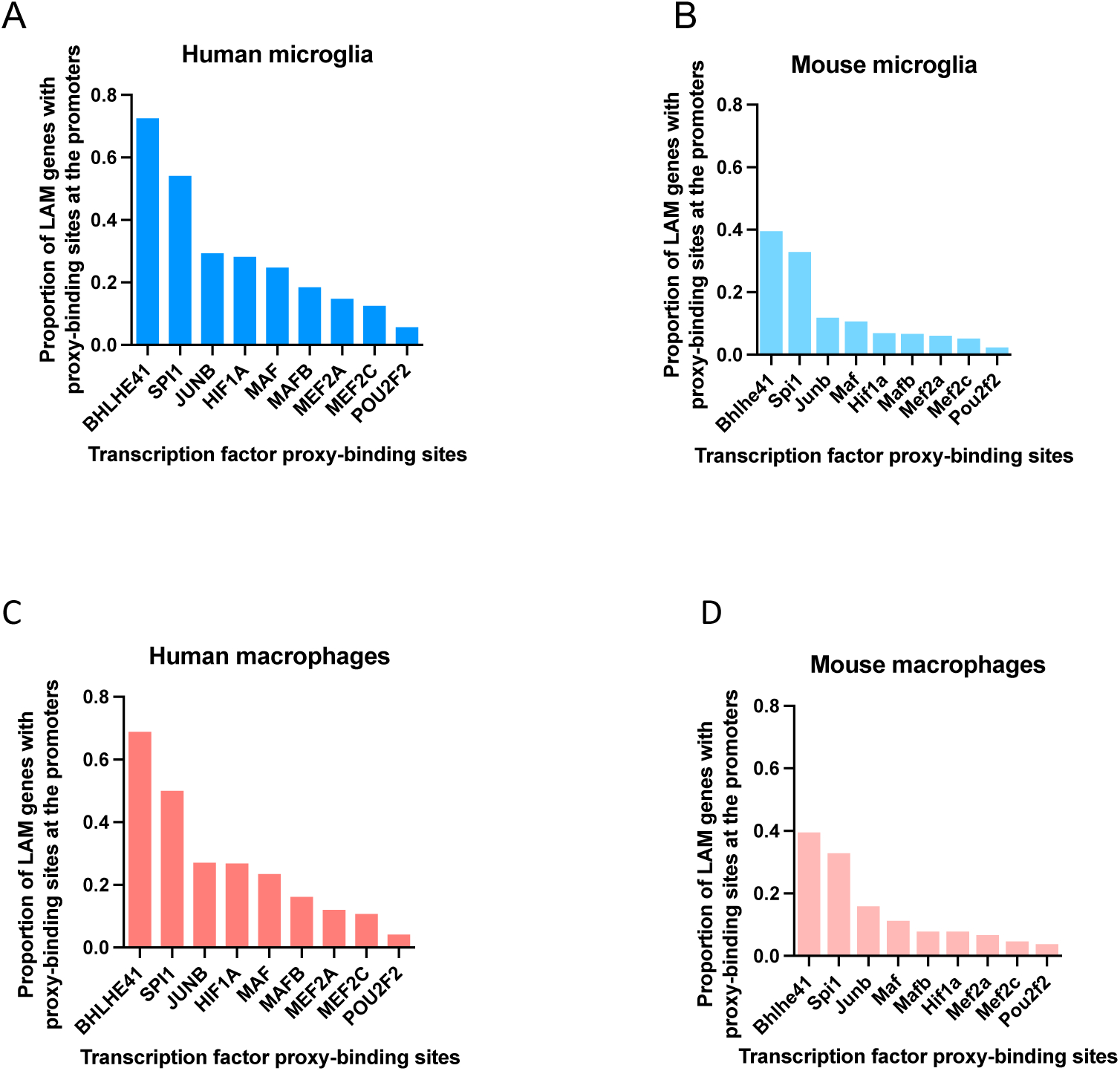
BHLHE41 and SPI1/PU.1 likely regulate lipid-associated macrophage (LAM) response genes through binding to their promoters. Proportions of LAM genes that contain TF proxy-binding sites in their promoter in **A)** human microglia, **B)** mouse microglia, **C)** human macrophages, **D)** mouse macrophages, for each candidate LAM TF listed on the x-axis.

### BHLHE40 and BHLHE41 are predicted to regulate LAM response genes involved in several pathways including cholesterol clearance and lysosomal processing

Since we and others have already been investigating the role of SPI1/PU.1 in microglia [33,34,46–48], we decided to focus our functional validation efforts for this study on the other strong candidate LAM TF BHLHE41 and its close homolog BHLHE40, which we (this study) and others [11,17,49] have nominated as a candidate transcriptional regulator of the LAM and similar neurodegeneration-related myeloid cell responses in mice, and which resides in the vicinity of a locus recently associated with AD risk in a GWAS of African-American individuals [38].

To investigate which of the LAM response-associated pathways are regulated by BHLHE40/41, we performed pathway analysis of 1) LAM genes with promoters containing BHLHE40 and/or BHLHE41 proxy-binding sites and 2) LAM genes belonging to the BHLHE40 and/or BHLHE41 regulons. We observed that BHLHE40 proxy-binding sites are a subset of BHLHE41 proxy-binding sites, thus we combined LAM response genes in (1) and refer to them as “LAM genes with promoters proxy-bound by BHLHE40/41” (Supplementary Table 2). Additionally, we observed a significant overlap between LAM genes in the BHLHE40 and BHLHE41 regulons (P-value = 1.59e-17). Hence, we combined LAM response genes in (2) and refer to them as “LAM genes in BHLHE40/41 regulons” (Supplementary Table 2).

To conduct pathway analysis we used Ingenuity Pathway Analysis (IPA, QIAGEN) [50] and identified pathways shared between “LAM genes with promoters proxy-bound by BHLHE40/41” and “LAM genes in BHLHE40/41 regulons”. They include “Phagosome maturation”, “Oxidative phosphorylation”, “Antigen presentation”, “Glycolysis”, “Atherosclerotic signaling”, “LXR/RXR activation”, and “mTOR signaling” (Supplementary Figure 2). Interestingly, these pathways are also regulated by other transcription factors such as LXR:RXR nuclear receptors and members of the MiT/TFE family TFEB and MITF, master regulators of cholesterol clearance and lysosomal processing, respectively [51]. These TFs are also known to directly target and stimulate the expression of BHLHE40/41 which in turn have been shown to act as repressors of transcriptional responses induced by LXR:RXR and MiT/TFE family TFs to form a negative feedback loop [39,40,52]. This suggests that regulation of the aforementioned LAM response-associated pathways is shared between known regulators of these pathways and BHLHE40/41.

### Knockout of BHLHE40/41 partially recapitulates the LAM transcriptional response in human iPSC-derived microglia (iMGLs)

To investigate the role of BHLHE40/41 as potential LAM TFs we generated an isogenic set of CRISPR-edited *BHLHE40* and *BHLHE41* single and double knockout (40KO, 41KO, and DKO) iPSC lines (Methods and Supplementary Figure 3A). Following microglial differentiation [53], we confirmed loss of BHLHE40 protein expression in 40KO and DKO iMGLs as well as loss of BHLHE41 protein expression in 41KO and DKO iMGLs by western blot (Supplementary Figure 3B).

Next, we performed RNA-seq analysis to determine the genes and pathways that are altered in 40KO, 41KO and DKO iMGLs compared to iMGLs derived from the parental (WT) iPSC line. We sequenced isogenic iMGLs of all four genotypes (WT, 40KO, 41KO and DKO) across five independent microglial differentiations for each genotype. Results of differential gene expression and gene set enrichment analyses are included in Supplementary Table 3.

Since each candidate LAM TF is predicted by our GRN analysis to strongly influence the expression of only a subset of LAM genes, we did not expect that transcriptional changes associated with loss of BHLHE40 and/or BHLHE41 in iMGLs *in vitro* could largely recapitulate the LAM transcriptional response observed in macrophages/microglia *in vivo*. Indeed, transcriptional changes in all KO iMGLs are significantly and positively but only modestly correlated with the LAM signature published by Jaitin *et al*. [54] (Dataset S6, FDR Adj.P-value < 0.05; Spearman’s correlation of ranks (ρ); ρ_40KO_ = 0.13 [0.04, 0.22], P-value_40KO_ = 0.003; ρ_41KO_ = 0.19 [0.1, 0.27], P-value_41KO_ <

0.0001; ρ_DKO_ = 0.30 [0.21, 0.37], P-value_DKO_ < 0.0001). Therefore, we used the rank-rank hypergeometric overlap (RRHO) analytical approach [55, 56] (Methods) to more precisely identify statistically significant overlaps between transcriptional changes associated with loss of BHLHE40 and/or BHLHE41 (KO vs WT) in iMGLs and the LAM signature. We used an improved stratified RRHO method [56] because this method can identify and visualize statistically significant overlaps between two transcriptional signatures regardless of whether they change in the same (concordant) or opposite (discordant) direction. This analytical approach revealed statistically significant and concordant overlaps in the bottom-left quadrant (i.e., between genes up-regulated in each pair of transcriptional signatures; hypergeometric overlap P-value: P-value_40KO_ = 1.90e-3, P-value_41KO_ = 2.28e-5, P-value_DKO_ = 2.74e-9) and in the top-right quadrant (i.e., between genes down-regulated in each pair of transcriptional signatures; hypergeometric overlap P-value: P-value_40KO_ = 5.30e-3, P-value_41KO_ = 5.38e-9, P-value_DKO_ = 1.40e-16) (Figure 4A and B). Interestingly, the most statistically significant and concordant overlaps were observed between the LAM signature and transcriptional changes that occur in DKO iMGLs (Figure 4A and B) suggesting that BHLHE40 and BHLHE41 may act together to regulate the expression of a subset of LAM response genes in human microglia. Next, We used IPA to perform pathway enrichment and activity analysis of RRHO overlapping up- and down-regulated genes from Figure 4B. Significantly enriched biological functions and diseases predicted to be more active (i.e., have positive Z-scores) in KO compared to WT iMGLs include “Endocytosis”, “Engulfment of cells”, “Transport of lipids”, “Migration of cells”, and “Antigen presentation”. Significantly enriched biological functions and diseases predicted to be less active (i.e., have negative Z-scores) in KO compared to WT iMGLs include “Accumulation of cholesterol”, “Obesity”, and “Lysosomal storage disease” (Figure 4C). Interestingly, these pathways point to increased activity of LXR:RXR and MiT/TFE family TFs which, as noted in the previous section, are master regulators of cholesterol clearance and lysosomal processing, respectively.

**Figure 4.**
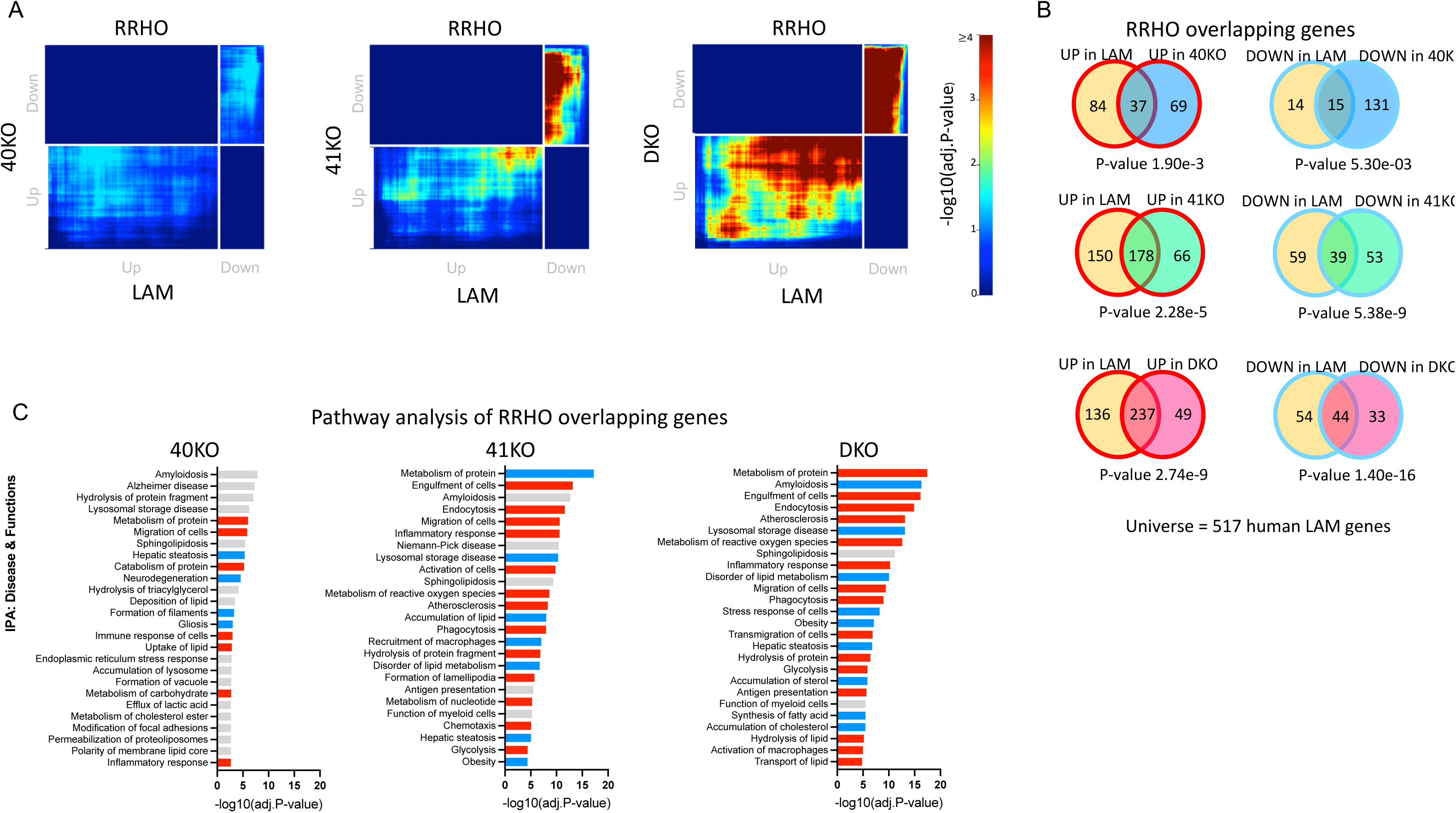
Knockout of *BHLHE40/41* partially recapitulates the LAM transcriptional response in human iPSC-derived microglia (iMGLs). **A)** Rank-rank hypergeometric overlap (RRHO) heatmaps visualizing significant overlaps in gene expression changes between each pair of BHLHE40/41 knockout (KO) iMGL and human LAM (Jaitin et al. [3], Dataset S6, FDR Adj.P-value < 0.05) transcriptional signatures. Adj.P-value in color temperature scale represents Benjamini-Hochberg corrected P-value of hypergeometric overlap test. B) Venn diagrams of most significant overlaps between genes up-regulated in both *BHLHE40/41* KO iMGL and human LAM transcriptional signatures, corresponding to the warmest pixel in the bottom-left quadrant of the respective heatmap (left) and Venn diagrams of most significant overlaps between genes down-regulated in both BHLHE40/41 KO iMGL and LAM transcriptional signatures, corresponding to the warmest pixel in the upper-right quadrant of the respective heatmap (right). P-values are calculated using the hypergeometric overlap test restricted to the universe of 517 human LAM genes (Jaitin *et al*. [3], Dataset S6, FDR Adj.P-value < 0.05). **C)** Pathways (“diseases and biological functions” category) found by IPA to be significantly enriched for RRHO overlapping genes. Red bars represent pathways predicted by IPA to be more active (i.e., with positive Z-scores), blue bars represent pathways predicted by IPA to be less active (i.e., with negative Z-scores), grey bars represent pathways with non-attributed Z-scores. Adj.P-value on the x-axis represents Bonferroni-Holm corrected P-value of pathway enrichment test. 40KO = BHLHE40 KO iMGLs, 41KO = BHLHE41 KO iMGLs, DKO = BHLHE40 and BHLHE41 double KO iMGLs, all compared to iMGLs derived from the parental iPSC line.

To further support these findings, we performed targeted differential gene expression analyses using RT-qPCR. We found that the expression of genes involved in cholesterol efflux/lipoprotein metabolism and lysosomal function was elevated in all KO iMGLs compared to WT iMGLs but reached statistical significance only in a few comparisons, likely due to low statistical power (Figure 5A, Supplementary Figure 4). Taken together these data suggest that loss of BHLHE40 and/or BHLHE41 induces the expression of LAM signature genes involved in cholesterol clearance and lysosomal processing.

**Figure 5.**
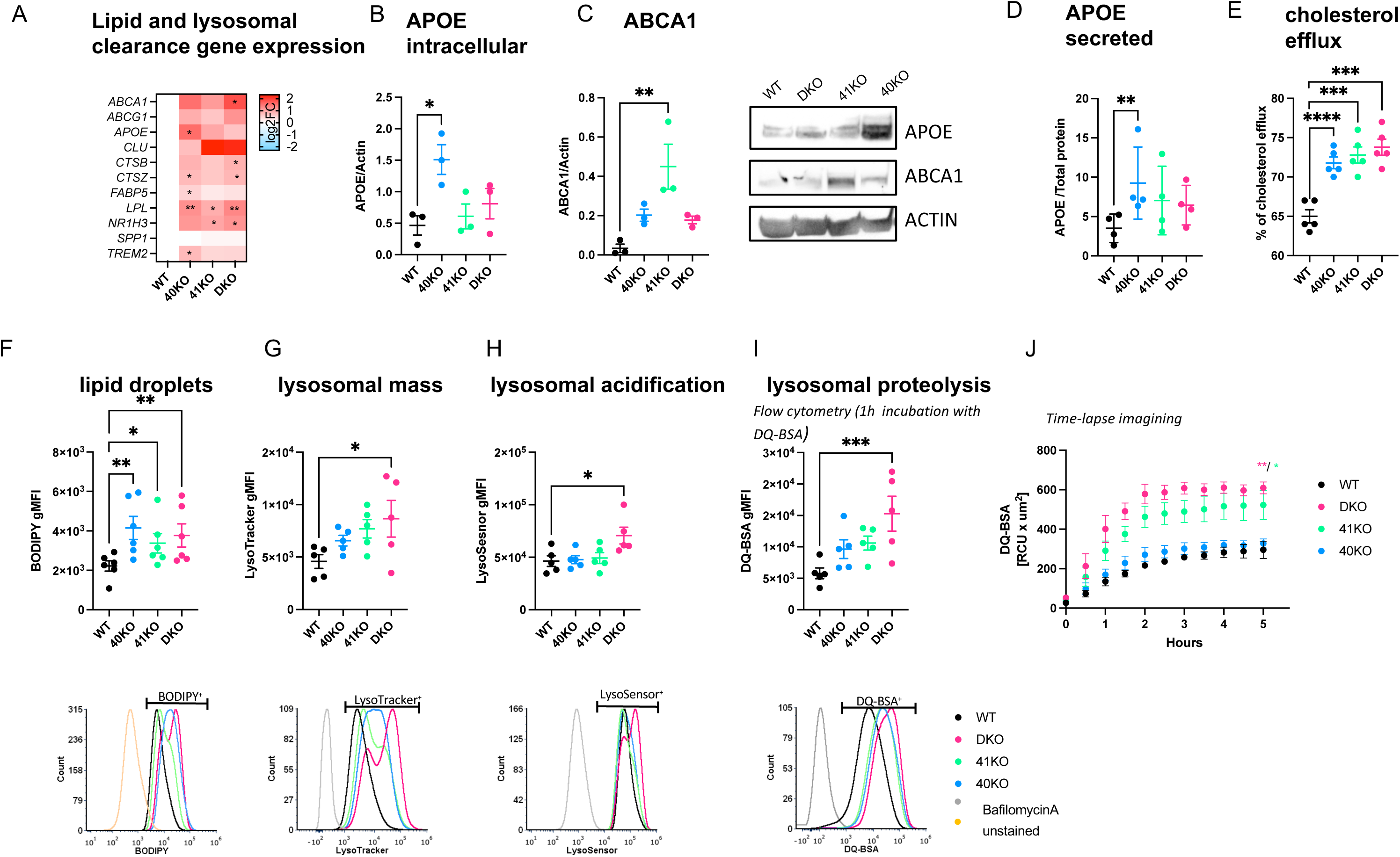
Knockout of *BHLHE40/41* increases expression of lipid and lysosomal clearance genes, cholesterol efflux and lipid droplet content, lysosomal mass and degradative capacity in human iPSC-derived microglia (iMGLs). **A)** Expression of lipid and lysosomal clearance genes measured by RT-qPCR, N=5/group. log2(fold change) (log2FC) is calculated with WT iMGLs as reference. Separate plots for each gene in Supplementary Figure 4. **B)** Intracellular APOE normalized to Actin measured by western blot, N=3/group. **C)** ABCA1 normalized to Actin measured by western blot, N=3/group. **D)** Secreted APOE normalized to total protein measured by ELISA, N=5/group. **E)** Percentage of cholesterol efflux, N=5/group. F) LDs content (BODIPY gMFI) measured by flow cytometry in BODIPY-positive cells (top), representative flow-cytometry histograms (bottom), N=6/group. **G)** Lysosomal mass (LysoTracker gMFI) measured by flow cytometry in LysoTracker-positive cells (top), representative flow-cytometry histograms (bottom), N=5/group. H) Lysosomal acidification (LysoSensor gMFI) measured by flow cytometry in LysoSensor-positive cells (top), representative flow-cytometry histograms (bottom), N=5/group. **I)** Lysosomal proteolysis (DQ-BSA gMFI) measured by flow cytometry in DQ-BSA-positive cells (top) representative flow- cytometry histograms (bottom), N=5/group. **J)** DQ-BSA red fluorescent signal (total integrated density) measured over time using the Incucyte S3 live imaging system, N=3/group. N represents the number of independent iMGLs differentiations. Flow cytometry gates were drawn based on FMO (fluorescence minus one) controls **(F)** or cells treated with Bafilomycin A **(G–I)** to prevent fusion of autophagosomes with lysosomes and thus obtain LysoSensor, LysoTracker and DQ-BSA negative signals. Differences of means between groups were tested using one-way ANOVA with repeated measures followed by Dunnett’s post-hoc test. * P-value < 0.05, ** P-value < 0.01, *** P-value < 0.001. Data plotted as mean ± SEM. Detailed statistics are shown in Supplementary File 1. 40KO = BHLHE40 KO iMGLs, 41KO = BHLHE41 KO iMGLs, DKO = BHLHE40 and BHLHE41 double KO iMGLs, WT = iMGLs derived from the parental iPSC line.

### Knockout of BHLHE40/41 increases cholesterol efflux and lipid droplet accumulation in iMGLs

Given that 1) loss of *BHLHE40/41* led to increased expression of LXR:RXR target genes (Figure 5A, Supplementary Figure 4), 2) pathway analysis indicated “LXR/RXR activation” is shared between BHLHE40/41 direct targets and LAM response genes (Supplementary Figure 2), and 3) BHLHE40/41 have been shown to repress LXR:RXR target genes (including cholesterol efflux genes *APOE, LPL, ABCA1, and ABCG1*) [39], we tested our KO and DKO iMGLs for LXR:RXR activation by measuring expression of LXR:RXR target genes *APOE* and *ABCA1*, as well as cholesterol efflux, in comparison to WT iMGLs. We first investigated whether the increased *APOE* and *ABCA1* mRNA levels that we observed using RT-qPCR (Figure 5A, Supplementary Figure 4) also resulted in increased protein levels. To this end, we performed western blot analysis and found significantly higher levels of APOE protein in 40KO iMGLs (effect size (ES) [95% confidence interval]; ES_40KO-WT_=1.04 [0.43, 1.68], P-value_40KO-WT_=0.0050, N=3/group), as well as elevated levels of ABCA1 in all KO iMGLs but reaching statistical significance only in 41KO iMGLs (ES_41KO-WT_=0.42 [0.21, 0.63], P-value_41KO-WT_=0.0021, N=3/group) (Figure 5B and C). We also observed significantly increased levels of secreted APOE in conditioned media from 40KO iMGLs cultures, consistent with the significantly higher levels of *APOE* mRNA and protein observed in these cells (Figure 5D) (ES_40KO-WT_=5.76 [2.03, 9.49], P-value_40KO-WT_=0.0048, N=5/group).

*APOE* and *ABCA1* are direct LXR:RXR target genes and are involved in cholesterol efflux and lipoprotein metabolism. Therefore, we measured the capacity of KO iMGLs to efflux cholesterol to an HDL acceptor. Cholesterol efflux capacity (measured as percentage of labeled cholesterol effluxed over 4h, Methods) was increased in all KO iMGLs compared to WT iMGLs (ES_40KO-WT_=6.80% [3.76%, 9.84%], P-value_40KO- WT_=0.0002; ES_41KO-WT_=7.80% [4.76%, 10.84%], P-value_41KO-WT_<0.0001; ES_DKO-WT_=8.80% [5.76%, 11.84%], P_DKO-WT_<0.0001; N=6/group), supporting our hypothesis that cholesterol clearance is enhanced by genetic inactivation of *BHLHE40* and/or *BHLHE41* in human microglia (Figure 5E).

Another feature of LXR:RXR activation that is also an attribute of LAMs is increased accumulation of lipid droplets (LDs) [3, 57]. Excess cholesterol delivered to macrophages by efferocytosis of lipid-rich cellular debris must be effluxed to extracellular acceptors like APOE or esterified for storage in intracellular LDs to prevent a cytotoxic buildup of free cholesterol [58]. To address the role of BHLHE40/41 in LDs accumulation, we took advantage of the fluorescent marker BODIPY which stains neutral lipids to measure LDs content by flow cytometry. We found that all KO iMGLs showed increased LDs content as evidenced by higher BODIPY geometric mean fluorescence intensity (gMFI) compared to WT iMGLs (ES_40KO-WT_=1928 [783.7, 3072], P-value_40KO-WT_=0.0014; ES_41KO-WT_=1151 [7.441, 2295], P-value_41KO-WT_=0.0484; ES_DKO-WT_=1546 [401.8, 2690], P-value_DKO-WT_=0.0082; N=6/group) (Figure 5F). Of note, LDs accumulation in DKO iMGLs is similar to that observed in alveolar macrophages of *Bhlhe40*/*41* double knockout mice [59]. To determine whether increased LDs content can be induced by LXR:RXR activation in iMGLs, we treated WT iMGLs with an LXR agonist (TO901317, 10µM for 48h) and observed higher BODIPY gMFI in these cells compared to WT iMGLs treated with vehicle as a control (Supplementary Figure 5A). Conversely, we treated KO iMGLs with an LXR antagonist (GW2033, 2µM, 24h) and saw that BODIPY gMFI in these cells was reduced to the levels observed in vehicle-treated WT iMGLs, indicating that LDs accumulation in *BHLHE40/41* KO iMGLs is caused by increased LXR:RXR activity (Supplementary Figure 5B).

A subpopulation of hippocampal microglia from aged mice accumulate LDs and display a pro-inflammatory phenotype distinct from LAMs [60]. To determine if increased LDs content in KO iMGLs is accompanied by increased secretion of pro-inflammatory cytokines, we measured levels of secreted cytokines in conditioned media. We found significantly lower levels of pro-inflammatory cytokines including MIF, IL13, CCL2 and IL1RA in all KO iMGLs compared to WT iMGLs (Supplementary Figure 6A-B), suggesting that the elevated LDs content in KO iMGLs is not associated with a pro-inflammatory phenotype. This observation is consistent with the suppression of pro-inflammatory cytokines by increased LXR:RXR activity that has been observed in peripheral macrophages that accumulate LDs (foam cells) [61].

### Knockout of BHLHE40/41 increases lysosomal mass and degradative capacity in iMGLs

RRHO overlapping genes between LAM and all KO iMGLs transcriptional signatures are enriched for pathways related to lysosomal function (Figure 4B and C), suggesting BHLHE40/41 may regulate lysosomal processing in human microglia. In addition, BHLHE40 and BHLHE41 have been shown to repress the master regulators of lysosomal biogenesis, TFEB and MITF, through binding to the same set of gene regulatory elements [40, 62]. These findings suggest that decreasing BHLHE40/41 expression may lead to enhanced activity of MiT/TFE family TFs and therefore increased lysosomal mass and degradative capacity. To test this hypothesis, we measured LysoTracker-Red and LysoSensor-Green gMFI by flow cytometry to quantify lysosomal mass and lysosomal pH, respectively. Consistent with our hypothesis, we observed an increase in lysosomal mass in all KO iMGLs, that reached statistical significance in DKO compared to WT iMGLs (ES_DKO-WT_=4080 [109.9, 8049], P-value_DKO-WT_=0.0438, N=5/group) (Figure 5G), as well as a statistically significant decrease in lysosomal pH (and therefore increased acidification of lysosomes) in DKO compared to WT iMGLs (ES_DKO-WT_=24256 [3502, 45009], P-value_DKO-WT_=0.0222; N=5/group) (Figure 5H). Lower lysosomal pH and increased expression of lysosomal cathepsins (Figure 5A) suggest enhanced degradative capacity of lysosomes in DKO iMGLs. To test this, we used a fluorochrome-conjugated Bovine Serum Albumin (DQ-BSA) assay. During endocytosis, DQ-BSA is delivered to the late endosome/lysosome and is subject to proteolysis by lysosomal enzymes leading to an increase in fluorescence. Using flow cytometry, we showed that BSA proteolysis (measured by gMFI of the conjugated fluorochrome) is significantly enhanced in DKO compared to WT iMGLs (ES_DKO-WT_=9485 [4678, 14293], P-value_DKO-WT_=0.0005, N=5/group) after 1h incubation with DQ-BSA (Figure 5I). Bigger differences between genotypes were observed during time-lapse recording (up to 5h, using Incucyte S3 live-cell imaging system, Figure 5J). We observed that DQ-BSA proteolysis is unchanged in 40KO iMGLs but it is increased in 41KO and DKO iMGLs compared to WT iMGLs (ES_41KO- WT_=192 [47.91, 336.0], P-value_41KO-WT_=0.0126; ES_DKO-WT_=281.6 [137.5, 425.6], P-value_DKO-WT_=0.0013, N=3/group) (Figure 5J). Altogether these data suggest that lysosomal function is increased in 41KO and DKO iMGLs compared to WT iMGLs.

### Knockdown of BHLHE40/41 partially recapitulates the LAM transcriptional and cellular response in human THP-1 macrophages

To further validate the role of BHLHE40/41 as modulators of the LAM response, we used the THP-1 model of human monocyte-derived macrophages (MACs). We transiently transfected MACs with siRNAs targeting *BHLHE40* (40KD), *BHLHE41* (41KD) or both (DKD). Using this approach, *BHLHE40* expression was reduced by ∼60% in *40*KD and DKD MACs, and *BHLHE41* expression was reduced by ∼50% in *41*KD and DKD MACs at both transcript and protein levels (Supplementary Figure 3B-C) when compared to control cells transfected with scrambled siRNA (SCR). Using THP-1 macrophages with transient reduction of *BHLHE40* and/or *BHLHE41* expression, we performed a similar set of experiments as we did in iMGLs with genetic inactivation of BHLHE40 and/or BHLHE41 (see previous sections) in order to validate our findings in an additional human macrophage cell culture model. First, we performed RNA-seq (N=4/group) to assess global transcriptomic changes associated with reduction of BHLHE40 and/or BHLHE41 expression in MACs. Results of differential gene expression and gene set enrichment analyses are included in Supplementary Table 4. Next, we used the RRHO method to identify statistically significant overlaps between the LAM signature published by Jaitin *et al*. [54] (Dataset S6, FDR Adj.P-value < 0.05) and transcriptional changes associated with reduction of BHLHE40 and/or BHLHE41 expression (KD vs SCR) in MACs. A significant overlap was identified between genes down-regulated in each pair of KD MAC and LAM transcriptional signatures (Supplementary Figure 7A) (hypergeometric overlap P-value: P-value_40KD_ = 1.44e-5, P-value_41KD_ = 7.96e-6, P-value_DKD_ = 3.13e-7) (Supplementary Figure 7B). A significant overlap was also identified between smaller subsets of genes up-regulated in 40KD or DKD MACs compared to SCR MACs and the LAM transcriptional signature (hypergeometric overlap P-value: P-value_40KD_ = 4.10e-3, P- value_DKD_ = 1.36e-6) (Supplementary Figure 7B). Similarly to what we observed using *in vitro* cultures of *BHLHE40/41* double-knockout iMGLs, the most statistically significant and concordant overlaps were observed between the LAM signature and transcriptional changes that occur in DKD MACs, suggesting that expression of both BHLHE40 and BHLHE41 have to be eliminated or reduced to more robustly mimic the LAM transcriptional response observed *in vivo*. Pathway enrichment and activity analysis of RRHO overlapping up- and down-regulated genes using IPA (Supplementary Figure 7C) revealed that “Efflux of cholesterol” and “Engulfment of cells” are predicted to be more active (i.e., have positive Z-scores) in KD compared to SCR MACs, and that “Lysosomal storage disease”, “Accumulation of lipid”, “Inflammatory response” are predicted to be less active (i.e., have negative Z-scores) in KD compared to SCR MACs (Supplementary Figure 7C). To further support these findings, we performed targeted differential gene expression analyses using RT-qPCR. We found that the expression of genes involved in cholesterol efflux/lipoprotein metabolism and lysosomal function was elevated especially in DKD MACs compared to SCR MACs but reached statistical significance only in a few comparisons, likely due to low statistical power and the fact that the extent of upregulation is smaller compared to that observed in iMGLs with complete loss of BHLHE40/41 expression (Figure 6A). To further investigate whether reduced expression of *BHLHE40* and/or *BHLHE41* in THP-1 macrophages could recapitulate the changes in cellular functions that we observed in iMGLs lacking *BHLHE40* and/or *BHLHE41*, we measured APOE secretion, cholesterol efflux capacity, and LDs content in KD and SCR MACs. We found that APOE secretion is significantly increased in *4*0KD compared to SCR MACs (Figure 6B) (ES_40KD-SCR_=2.33 [0.04, 4.62], P-value_40KD-SCR_=0.0457, N=6/group). We also found that cholesterol efflux to APOA-1 acceptor is significantly increased in all KD MACs compared to SCR MACs (ES_40KD-SCR_=12.45 [8.47, 16.44], P_40KD-SCR_<0.0001; ES_41KD- SCR_=4.83 [0.84, 8.81], P_41KD-SCR_<0.0170; ES_DKD-SCR_=6.91 [2.92, 10.89], P-value_DKD-SCR_=0.0011; N=6/group) (Figure 6C). Furthermore, we found that all KD MACs showed increased LDs content as evidenced by higher BODIPY gMFI compared to SCR MACs (ES_40KD-SCR_=2348 [567, 4128], P-value_40KD-SCR_=0.0097; ES_41KD-SCR_=2890 [1110, 4671], P-value_41KD-SCR_=0.0020; ES_DKD-SCR_=3563 [1782, 5343], P-value_DKD-SCR_=0.0003; N=6/group). We also determined that higher LDs content is not associated with a pro-inflammatory phenotype, as evidenced by reduced levels of pro-inflammatory cytokines secreted by all KD compared to SCR MACs (Supplementary Figure 6C-D).

**Figure 6.**
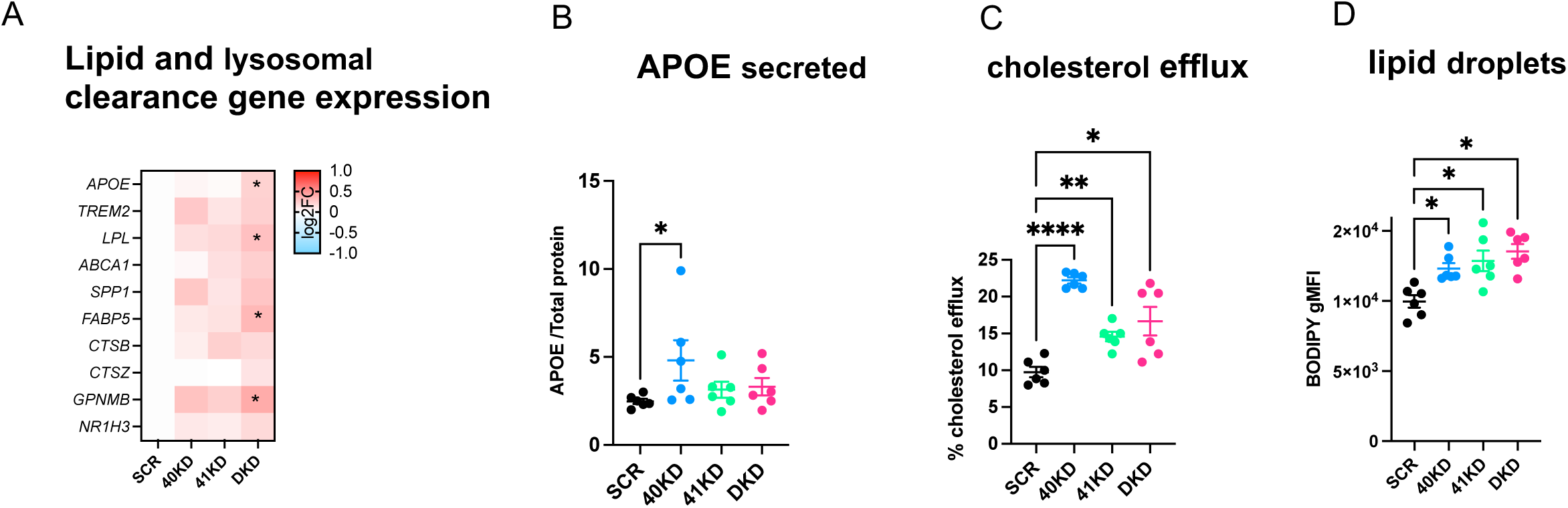
Knockdown of *BHLHE40/41* increases expression of lipid clearance genes, cholesterol efflux and lipid droplet content in human THP1 macrophages (MACs). **A)** Expression of lipid and lysosomal clearance genes measured by RT-qPCR, N=7/group. log2(fold change) (log2FC) is calculated with SCR MACs as reference. **B)** Secreted APOE normalized to total protein measured by ELISA, N=6/group. **C)** Percentage of cholesterol efflux, N=6/group. **D)** LD content (BODIPY gMFI) measured by flow cytometry in BODIPY-positive cells, N=6/group. N represents the number of independent siRNA transfections. Flow cytometry gates were drawn based on FMO (fluorescence minus one) controls. Differences of means between groups were tested using one-way ANOVA with repeated measures followed by Dunnett’s post-hoc test. * P-value < 0.05, ** P-value < 0.01, **** P-value < 0.0001. Data plotted as mean ± SEM. Detailed statistics are shown in Supplementary File 1. 40KD = MACs treated with BHLHE40 siRNA, 41KD = MACs treated with BHLHE41 siRNA, DKD = MACs treated with BHLHE40 and BHLHE41 siRNA, SCR = MACs treated with scrambled siRNA.

Together these data further demonstrate a novel role of BHLHE40/41 in the modulation of LAM response genes and cellular processes involved in lipid metabolism in another *in vitro* model of human macrophages and reveal a high degree of similarity between iMGLs with complete loss of BHLHE40/41 and THP-1 macrophages with reduced levels of *BHLHE40/41*.

### Knockout of Bhlhe40 and Bhlhe41 partially recapitulate the LAM transcriptional response in mouse microglia

To investigate whether loss of BHLHE40/41 function could lead to gene expression changes that partially recapitulate the LAM transcriptional response across species as well as *in vivo*, we acutely isolated microglia from *Bhlhe40*/*Bhlhe41* double knockout (DKO) mice [59] and their WT littermates and performed bulk RNA-seq transcriptome profiling. Results of differential gene expression and gene set enrichment analyses are included in Supplementary Table 5. Similarly to what we observed using *in vitro* cultures of *BHLHE40/41* double-knockout human iPSC-derived microglia, transcriptional changes in *Bhlhe40/41* double-knockout mouse microglia are significantly and positively correlated !"_DKO_ = 0.31, P-value_DKO_ < 0.0001) with the LAM signature published by Keren-Shaul *et al*. [10] (Table S3 FDR Adj.P-value < 0.05). Next, we used the RRHO method to identify statistically significant overlaps between the LAM signature and transcriptional changes associated with inactivation of both *Bhlhe40* and *Bhlhe41* (DKO vs WT) in mouse microglia. This analytical approach revealed statistically significant and concordant overlaps in the bottom-left quadrant (i.e., between genes up-regulated in both transcriptional signatures) (hypergeometric overlap P-value: P-value_DKO_ = 9.30e-18) and in the top-right quadrant (i.e., between genes down-regulated in both transcriptional signatures) (hypergeometric overlap P-value: P-value_DKO_ = 8.50e-24) (Figure 7A). Pathway enrichment and activity analysis of RRHO overlapping up- and down-regulated genes using IPA (Figure 7B) revealed that “Endocytosis”, “Phagocytosis”, and “Fatty acid metabolism” are predicted to be more active (i.e., have positive Z-scores) in DKD compared to WT mouse microglia, and that “Lysosomal storage disease”, “Gliosis”, and “Amyloidosis” are predicted to be less active (i.e., have negative Z-scores) in DKD compared to WT mouse microglia (Figure 7C). Taken together, these results indicate that transcriptional changes associated with loss of *Bhlhe40* and *Bhlhe41* in mouse microglia can partially recapitulate the LAM transcriptional response *in vivo*.

**Figure 7.**
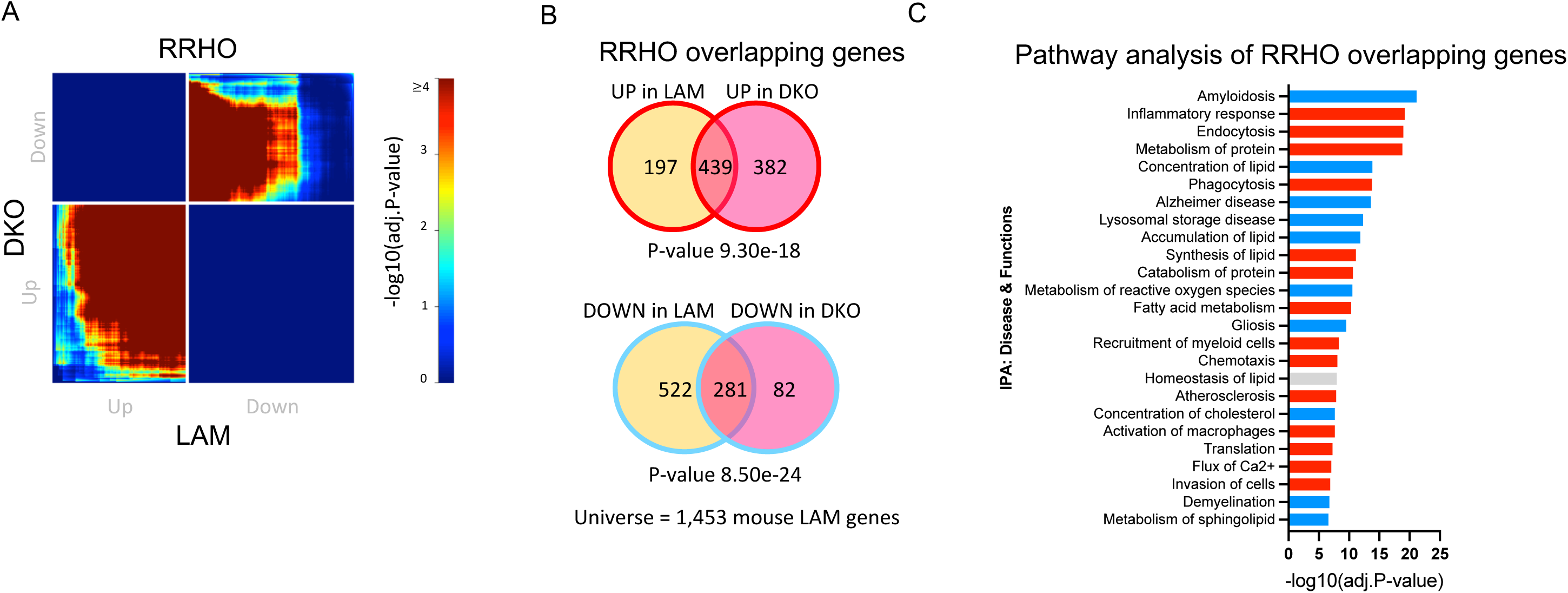
Knockout of *Bhlhe40* and *Bhlhe41* partially recapitulate the LAM transcriptional response in mouse microglia. **A)** Rank- rank hypergeometric overlap (RRHO) heatmaps visualizing significant overlaps in gene expression changes between mouse *Bhlhe40* and *Bhlhe41* double knockout (DKO) mouse microglia and mouse LAM (Keren-Shaul *et al*. [10], Table S3, FDR Adj.P-value < 0.05) transcriptional signatures. Adj.P-value in color temperature scale represents Benjamini-Hochberg corrected P-value of hypergeometric overlap test. **B)** Venn diagram of the most significant overlap between genes up-regulated in both *Bhlhe40/41* DKO mouse microglia and the mouse LAM transcriptional signature, corresponding to the warmest pixel in the bottom-left quadrant of the heatmap (top) and Venn diagram of the most significant overlap between genes down-regulated in both *Bhlhe40/41* DKO mouse microglia and the mouse LAM transcriptional signature, corresponding to the warmest pixel in the upper-right quadrant of the heatmap (bottom). P-values are calculated using the hypergeometric overlap test restricted to the universe of 1,453 mouse LAM genes (Keren-Shaul *et al*. [10], Table S3, FDR Adj.P- value < 0.05). **C)** Pathways (“diseases and biological functions” category) found by IPA to be significantly enriched for RRHO overlapping genes. Red bars represent pathways predicted by IPA to be more active (i.e., with positive Z-scores), blue bars represent pathways predicted by IPA to be less active (i.e., with negative Z-scores), grey bars represent pathways with non-attributed Z-scores. Adj.P-value on the x-axis represents Bonferroni-Holm corrected P-value of pathway enrichment test. DKO = *Bhlhe40/41* DKO mouse microglia, compared to microglia derived from wild-type control mice.

## Discussion

Macrophages that reside in cholesterol/lipid-rich tissues display similar TREM2- dependent transcriptional responses to host tissue damage (e.g., disease-associated microglia in the aging or degenerating brain, the most cholesterol/lipid-rich organ of the human body) [3,12,18–29]. We collectively refer to macrophages in this transcriptional state as lipid associated macrophages (LAM). In this study we report the identification of transcriptional regulators of the LAM response (LAM TFs) through gene regulatory network analyses of macrophage transcriptomes from human and mouse single-nucleus and single-cell RNA-seq data. In particular, we leveraged data not only from microglia in healthy and diseased brains, but also from macrophages in atherosclerotic plaques and adipose tissue from healthy and obese animals as well Kupffer cells in healthy and diseased livers [3,12,18–29]. Taking advantage of the breadth of challenge conditions in mouse models and human diseases as well as the high resolution of single-cell and single-nucleus RNA-seq data, we were able to nominate 11 strong candidate LAM TFs (*BHLHE41*, *HIF1A*, *ID2*, *JUNB*, *MAF*, *MAFB*, *MEF2A*, *MEF2C*, *NACA, POU2F2* and *SPI1*) that were highly replicated within and across species and conditions.

### The LAM response is likely regulated by cooperation of multiple TFs

Interestingly, several of these LAM TFs have been shown to interact with each other in various ways, suggesting that they might regulate the LAM response cooperatively. For example, PU.1 binds an enhancer element located downstream of the *Mef2c* hematopoietic-specific promoter and is required for regulating *Mef2c* expression during myeloid cell development [63]. Similarly, Bhlhe40 acts together with PU.1 to drive the selection of enhancers in large peritoneal macrophages in the presence of retinoic acid [64]. These data suggest that the candidate LAM TFs nominated in this study likely interact with each other in complex ways to cooperatively regulate the transcriptional responses of macrophages/microglia to cholesterol/lipid-rich tissues damaged by aging and disease.

Of note, several LAM genes in both human and mouse networks are predicted to be strongly influenced by only a small fraction of candidate LAM TFs. There are multiple reasons why this could be the case. Genuine biological specificity might be one of the reasons for this observation. Additionally, if the genes are very highly specific to the LAM response (e.g, *LPL*), the presence of only a small fraction of cells that significantly upregulate this marker in human tissues might make it difficult to infer a robust association with a TF. Furthermore, genes that are lowly expressed but significantly induced in LAM may not pass the expression threshold in our analyses for a proportion of the networks and thus do not display more associations with candidate TFs. Finally, robust changes in mRNA levels are required to infer associations between a gene and a TF with our approach; hence, for those genes whose mRNA values are not robustly varied, our approach would be unable to detect TF-gene associations.

### The role of LAM TFs and their upstream regulators in AD risk modification

We next sought to identify which of the nominated LAM TFs are more likely to directly regulate the LAM response, e.g., through binding to promoters of LAM genes. To achieve this, we used open chromatin profiles of macrophages/microglia and sequence motifs to which each TF is predicted to bind in order to identify TF proxy-binding sites, i.e., open chromatin regions (ATAC-seq peaks) that contain binding motifs for a specific TF. The highest proportion of LAM gene promoters contained a *BHLHE40/41* proxy-binding site in human and mouse macrophages/microglia, highlighting these two highly homologous and functionally redundant TFs as the most likely direct regulators of the LAM response. The interplay between AD risk genes and regulation of the LAM response has been previously highlighted [65], suggesting that AD risk genes may modulate disease susceptibility by regulating the gene expression profiles and in turn the cellular activities of macrophages in response to dyshomeostasis of cholesterol/lipid-rich tissues [8, 9]. Consistent with this hypothesis, previous studies have reported that several well-established AD risk genes (*APOE*, *TREM2* and *PLCG2*) are critically important for the development and function of LAM in response to cholesterol/lipid overload during aging and disease [10,11,66]. Furthermore, a recent study showed that APOE risk-increasing (*APOE ε4/ε4*, similar to *APOE*, *TREM2*, and *PLCG2* loss-of-function mutations) and risk-decreasing (*APOE ε2/ε2*) genotypes are associated with decreased and increased LAM transcriptional response (as well as decreased and increased efferocytosis), respectively, compared to the risk-neutral APOE *ε3/ε3* genotype [67]. Additionally, several TFs that we nominated in this study as candidate LAM TFs (*MAF, MEF2C*, and *SPI1*, encoding the myeloid master regulator PU.1) were also nominated as candidate AD risk genes in genome-wide association studies (GWAS) [33,68,69]. We and others have also previously shown that AD heritability is enriched within PU.1 (proxy- as well as ChIP-seq) binding sites in macrophages/microglia and that AD risk alleles in the *SPI1* AD GWAS locus affect PU.1 expression, further implicating this LAM TF in the modulation of AD risk [33,36,70]. Here we show that AD heritability is similarly enriched within proxy-binding sites of BHLHE41 (and its close homolog BHLHE40) in macrophages/microglia, implicating the BHLHE40/41 target gene network not only in LAM response regulation but also in AD risk modification. Interestingly, *BHLHE40* is located near an AD locus recently identified in an African-American GWAS [38] and therefore should be considered as a candidate AD risk gene for future studies in macrophages/microglia derived from this human population.

### Putative mechanism of BHLHE40/41-mediated regulation of the LAM response

*Bhlhe40* is upregulated in mouse LAM and has been proposed to regulate the expression of genes in neurodegeneration- and demyelination-related co-expression modules in mouse microglia [11,17,49,71]. However, the role of *BHLHE40* (and its close homolog *BHLHE41*) in the regulation of LAM genes and pathways has not been experimentally validated or further investigated. In the current study, we used human iPSC-derived microglia and THP-1 macrophages to show that complete loss or reduced levels of BHLHE40 and/or BHLHE41 in these cells is associated with enhanced expression of LAM genes and increased activity of LAM pathways (i.e., cholesterol clearance and lysosomal processing) suggesting that these TFs, when induced during the LAM response, may act as transcriptional repressors of these functional components of the LAM response. Inhibition of the LAM response by BHLHE40/41 may be part of a negative feedback loop initiated by other TFs such as LXR:RXR (and other RXR-containing) nuclear receptors and MiT/TFE family members [39,40,72]. Indeed, studies show that LXR:RXR activation induces expression of *Bhlhe40* [73] which in turn (together with Bhlhe41) represses expression of genes induced by LXR:RXR activation [39]. Similar repression by *Bhlhe40* have been shown for other TFs such as PPARγ:RXRɑ whose target genes are also involved in lipid metabolism (i.e. lipid storage, lipolysis) and are markedly increased in *Bhlhe40* deficient mice in white adipose tissue and liver [41, 74]. Similarly, *BHLHE40* and *BHLHE41* gene expression is induced by nuclear TFEB and act in opposition to TFEB, competing for the same binding sites [40]. Another example is MITF, which shares similar functions with TFEB and promotes the expression of genes involved in lysosomal biogenesis. In melanoma, *BHLHE40* and *BHLHE41* transcription is induced by direct binding of MITF to their promoters [52]. In turn, MITF transcription is repressed by direct binding of BHLHE40 to the *MITF* promoter [62]. Consistent with these observations, the results of our experiments using human iPSC-derived microglia and THP-1 macrophages indicate that interrupting/weakening this negative feedback loop by eliminating/reducing *BHLHE40* and/or *BHLHE41* gene expression in these cells is associated with increased expression of select LXR:RXR and MiT/TFE target genes and with increased activity of pathways regulated by these TFs, such as cholesterol clearance and lysosomal processing (Figure 8). This model implies that LXR:RXR nuclear receptors and MiT/TFE family members may themselves be LAM TFs. Indeed, MITF has been nominated and functionally validated as a candidate driver of a LAM-like transcriptional state observed in cultured iMGLs exposed to cholesterol/lipid-rich cellular debris *in vitro* [32]. In addition, LXRα/β have been nominated as candidate drivers of an anti-inflammatory LAM transcriptional state observed in microglia acutely isolated from the brain of 5XFAD mice [15].

**Figure 8.**
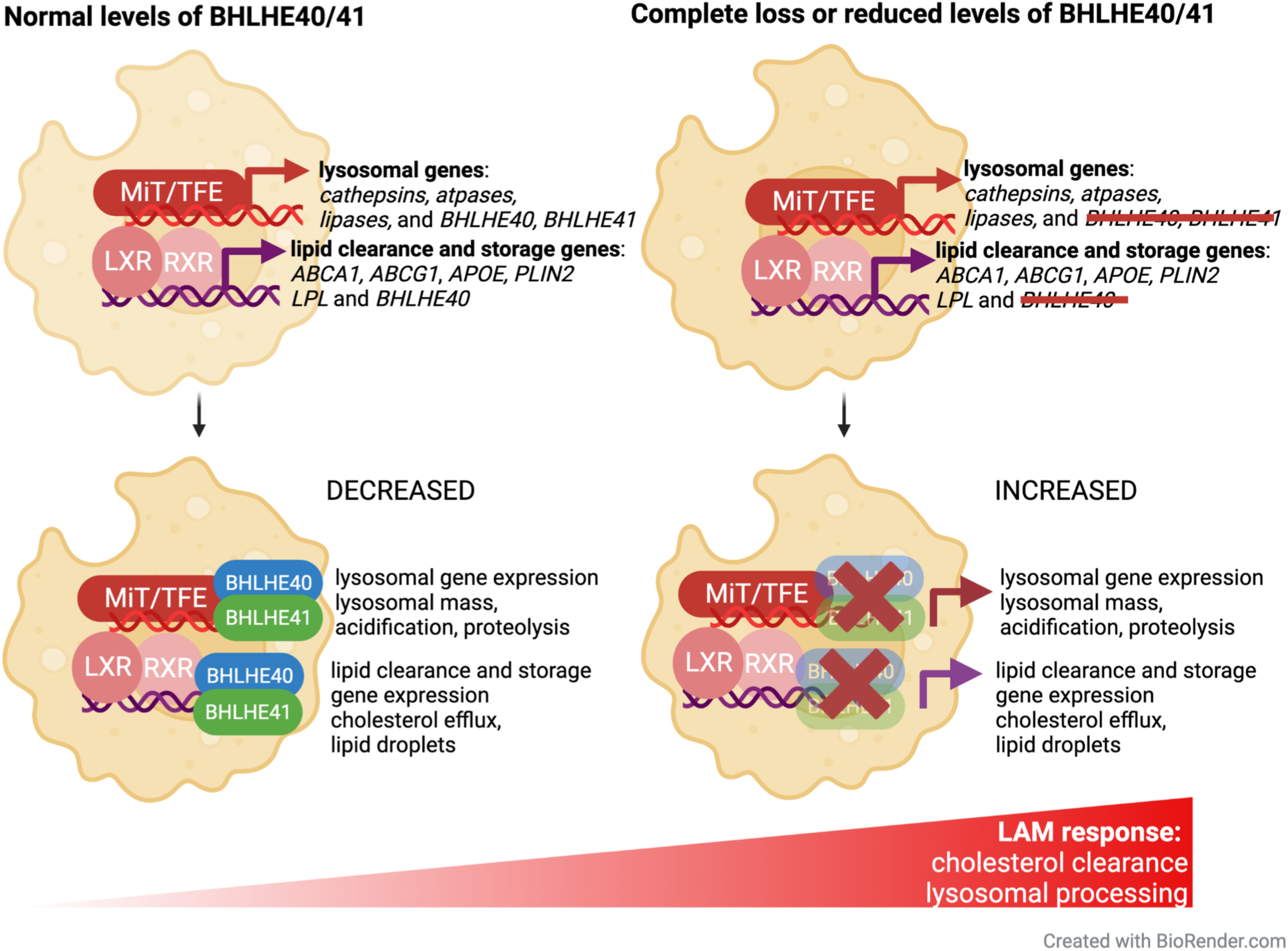
Proposed mechanism for LAM response regulation by BHLHE40/41. **Left)** In wild-type mouse microglia or human iPSC-derived microglia (iMGLs) and THP-1 macrophages (MACs) treated with scrambled siRNA (and thus with normal levels of BHLHE40/41), when LXR:RXR and MiT/TFE family TFs stimulate the expression of cholesterol clearance and lysosomal processing genes that contribute to the LAM transcriptional program, they also stimulate the expression of *BHLHE40* and *BHLHE41*. In turn, BHLHE40/41 oppose the activity of LXR:RXR and MiT/TFE family TFs by binding to the promoters of their target genes and leading to decreased expression of cholesterol clearance and lysosomal processing genes, in a negative feedback loop. **Right)** In mouse microglia or human iMGLs and MACs with complete loss or reduced levels of BHLHE40/41, this negative feedback loop is interrupted or weakened resulting in increased expression of cholesterol clearance and lysosomal processing genes to partially recapitulate the LAM transcriptional and cellular response, including increased cholesterol efflux to APOE and storage in LDs, as well as increased lysosomal mass, acidification and proteolysis.

### Lipid droplet phenotype associated with BHLHE40/41 loss-of-function in macrophages/microglia

In this study, we show that complete or partial loss of BHLHE40/41 function in human iPSC-derived microglia (iMGLs) and THP-1 macrophages (MACs) cultured *in vitro* is associated with increased levels of neutral lipids/lipid droplets (LDs), a phenotype that we have also shown to be dependent on LXR:RXR activation. Increased LDs content has also been observed *in vivo* in alveolar macrophages of *Bhlhe40*/*41* double knockout mice [59]. These findings are in accordance with recent studies reporting a similar LDs phenotype in LAM populations of human and mouse macrophages/microglia [3, 57]. For example, a subset of macrophages found in adipose tissue of obese individuals and characterized by high levels of LAM markers such as CD9 and LPL also show increased LDs accumulation [3]. In the context of AD, a recent study showed that plaque-associated xenotransplanted microglia (xMGLs, human iPSC-derived microglia injected into the brain of immunocompetent mice [75]) acquire a LAM transcriptional signature and accumulate PLIN2 (perilipin 2)-positive LDs [57]. In contrast, other studies identified a subset of microglia that accumulate LDs in the hippocampus of aged mice (referred to by the authors as LAM, for LD-associated microglia) and are characterized by increased expression of pro-inflammatory genes and secretion of pro-inflammatory mediators [76]. In *BHLHE40/41* KO/KD iMGLs and MACs, increased LDs content is, on the contrary, associated with lower levels of pro-inflammatory mediators secreted in the culture media. These data suggest that increased accumulation of LDs due to lack of *Bhlhe40/41* is not associated with a pro-inflammatory phenotype and may instead be part of a protective adaptive response in which sequestration of free cholesterol and other potentially toxic lipids prevents cellular stress and inflammation [77–80]. Interestingly, pharmacological activation of LXR:RXR nuclear receptors leads to an increase in LDs in human retinal pigmented epithelial cells and promotes both cholesterol efflux and LDs formation [81] similar to what we have observed in *BHLHE40/41* KO/KD iMGLs and MACs. This suggests that LXR:RXR activation downstream of BHLHE40/41 loss-of-function in macrophages/microglia promotes cholesterol clearance by stimulating both its efflux to extracellular acceptors like APOE and its esterification for non-toxic accumulation and intracellular storage in LDs [81]. All together, these findings support our hypothesis that the higher LDs content observed in macrophages/microglia in the absence of BHLHE40/41 may be attributed to increased LXR:RXR activity in these cells and may be a part of a protective mechanism preventing cellular stress and inflammation induced by potentially toxic levels of lipids like free cholesterol.

### Study limitations

Although we were able to pinpoint and functionally validate transcriptional regulators of the LAM response, there are several limitations to our study. The LAM signature has been robustly identified in several mouse models of aging and disease, however its identification in human tissues has been more challenging [82]. One of the challenges is the prevalent use of snRNA-seq to identify human microglial transcriptional states [83]. On the other hand, a recent preprint study using scRNA-seq identified a transcriptional state in human iPSC-derived microglia cultured *in vitro* and treated with myelin fragments, apoptotic cells and synaptosomes that resembles the LAM signature observed in mouse microglia *in vivo*, suggesting an evolutionarily conserved pattern of microglial gene expression changes in response to cholesterol/lipid-rich cellular waste [32]. In this study we assume conservation of the LAM signature from mouse to human; however, as more single-nucleus and single-cell transcriptomic and multi-omic datasets from the aging and diseased human brain are generated, our understanding of the LAM signature in humans may evolve. Additionally, our analysis of microglial gene expression from double knockout of *Bhlhe40/41* shows that the LAM program is partially recapitulated, including pathways involved in “Lipid transport”, “Oxidative phosphorylation” and “Phagosome maturation”. These analyses were conducted on relatively young (2 and 6 month-old) and healthy mice; however, in future studies we will examine the LAM response in older mice or in the context of a disease-relevant challenge (e.g. mouse models of demyelination or amyloid deposition). Finally, we have utilized open chromatin data to generate sets of proxy- binding sites by stratifying ATAC-seq peaks based on the presence of known TF binding motifs. Since these motifs can be quite similar for related TFs, our proxy-binding sites likely reflect putative binding of multiple TFs within the same family. As more TF-specific ChIP-seq datasets from human and mouse microglia are generated, we will be able to further validate and refine our findings based on proxy-binding sites.

In summary, we have utilized single-cell and single-nucleus RNA-seq data from various human and mouse macrophage populations that were shown to exhibit a shared transcriptional response to cholesterol/lipid overload. We implicated multiple candidate transcriptional regulators of this response, with *BHLHE40/41* being the most likely direct regulators. We further implicated *BHLHE40/41* in AD risk modification, highlighting an intriguing interplay between transcriptional regulation of the LAM response and modulation of disease susceptibility. Finally, we validated these bioinformatic findings experimentally in human and mouse macrophages/microglia, nominating BHLHE40/41 as candidate transcriptional regulators of the LAM response and putative drug targets for therapeutic modulation of macrophage/microglial function in AD and other disorders of lipid-rich tissues.

## Methods

### Reconstruction of gene regulatory networks

Single-cell and single-nucleus RNA-seq datasets from microglia, Kupffer cells, atherosclerotic plaque macrophages, and adipose tissue macrophages were obtained from Gene Expression Omnibus (GEO) [3,12,18–29] (Supplementary Table 1). Cell type annotations reported by the authors were used to extract microglia, Kupffer cells, and other macrophages. Count matrices were obtained from each study and CPM normalized. Pearson correlation matrices were used as distance matrices for metacell reconstruction. The pipeline for reconstruction of metacells can be found at the accompanying website [84]. All functions were used with their default parameters. Briefly, this approach constructs a kNN graph using the data, partitions the data into an appropriate number of metacells, taking into account the desired number of neighbors. It then aggregates the counts from closest neighbors into MetaCells. The MetaCells function was used in the following manner: *MetaCells(data, dist.mat, numNeighbors = numNeighbors, subSize = subSize)*. Resulting metacells were filtered to include genes that showed non-zero counts in at least 75% of metacells. ARACNe was then used to reconstruct gene regulatory networks using default parameters [30]. The threshold for MI was calculated as follows: *java -Xmx5G -jar Aracne.jar -e Matrix_file.txt -o . --tfs list_of_TFs.txt --pvalue 1E-8 --seed 1 --calculateThreshold*. The network was built using 200 bootstraps using the following command for each: java -Xmx5G -jar Aracne.jar -e *Matrix_file.txt* -o . --tfs *list_of_TFs.txt* --pvalue 1E-8 --seed *seed_number*. Bootstraps were further consolidated into a single network in the following manner: *java -Xmx5G -jar Aracne.jar -o . --consolidate*. Only significant interactions (FDR Adj.P-value < 0.05) were used for further downstream analyses. Meta-analyzed networks (displayed in Figure 1A and B and used for final TF regulon extraction) were generated by aggregating all the bootstraps from individual networks generated by ARACNe using the *consolidate* function mentioned above [30, 84].

### Enrichment analyses in TF regulons

Each TF regulon was tested for significant overlap with three LAM genesets (human or mouse DAM, LAM, and TREM2^hi^ genesets, Supplementary Table 1) by using the hypergeometric test (*phyper* function in R *stats* package). The TFs whose regulons were significantly enriched (FDR Adj.P-value < 0.2) were nominated as LAM TFs. The LAM genesets for the enrichment analyses were obtained from their respective studies in the following manner: mouse DAM geneset was obtained from Table S2 of Keren-Shaul *et al*. [10], mouse LAM geneset was obtained from Dataset S5 of Jaitin *et al*. [3], and mouse TREM2^hi^ macrophage geneset was obtained from Online Table I (Cluster 11) of Cochain *et al*. [26]. The human LAM geneset was obtained from Dataset S6 of Jaitin *et al*. [3] and filtered for FDR Adj.P-value < 0.05. The human DAM and TREM2^hi^ genesets were constructed by obtaining the human orthologs of the mouse genes in the corresponding mouse geneset using biomaRt [85]. High confidence TFs were selected if they were 1) enriched for all three LAM genesets in at least half of human and mouse networks. 2) conserved between species, and 3) expressed (TPM ≥ 1) in human microgila [31].

### Processing of ATAC-seq data and peak calling

Macrophage ATAC-seq data were obtained through the Gene Expression Omnibus (GEO, GSE100383) [86]. Human microglial ATAC-seq data were obtained from the database of Genotypes and Phenotypes (dbGaP, phs001373.v2.p2) [31]. Mouse macrophage and microglia ATAC-seq data were obtained through GEO (GSE151015, GSE89960) [31, 87]. To generate epigenomic annotations, FASTQ files were obtained from the Sequence Read Archive (SRA) [88]. Technical replicates were merged and Bowtie2 was used for alignment for both single and paired-end files [89]. FASTQC was used for quality control of the files [90]. Resulting SAM files were filtered by MAPQ score and duplicates were removed using samtools [91]. MACS2 was used to call peaks from ATAC-seq data [92].

### Generation of promoter annotations and overlap with TF proxy-binding sites

We obtained TSS positions of genes using biomaRt [85] and extended them by 2kb on each side of the TSS to generate gene promoter annotations. To assess if a TF is likely to be bound at a gene promoter, we created *GRanges* objects for both promoter and open chromatin regions (ATAC-seq peaks) and utilized the *subsetByOverlaps* function from the *GenomicRanges* package [93] to identify open chromatin regions that contain binding motifs for the TF of interest (TF proxy-binding sites) and overlap with the gene promoter.

### De novo TF binding motif discovery and TF binding motif frequency analysis

We used HOMER to quantify the frequencies of TF binding motif occurrence in open chromatin regions that overlap gene promoter annotations [37]. The following command was used to quantify motif frequencies depicted in Figure 3: *annotatePeaks.pl Peaks.bed hg19 -m TF.motif -size 2000 -hist 20*. To identify open chromatin regions that contain binding motifs for the TF of interest, we used the following commands: *findMotifsGenome.pl Peaks.bed hg19 . -find motif.motif -size given* and *annotatePeaks.pl Peaks.bed hg19 -m motif.motif -size given*.

### Partitioned SNP heritability analysis

We used LD Score regression [94] to estimate AD SNP heritability partitioned by proxy- binding sites of candidate LAM TFs. We used GWAS summary statistics (excluding the APOE [chr19:45000000–45800000] and MHC/HLA [chr6:28477797–33448354] regions) from the Lambert *et al.* AD GWAS study [68].

### Generation of human BHLHE40 and/or BHLHE41 knockout iPSC lines

BHLHE40 and BHLHE41 single homozygous knock-out iPSC lines were generated by CRISPR/Cas9-mediated homologous direct repair (HDR). For BHLHE40 the HDR was designed to introduce p.R67>X, a premature stop codon in the bHLH domain. For BHLHE41 the HDR was designed to introduce p.T41>X, a premature stop codon in the bHLH domain. Briefly, one million human iPSCs from the WTC-11 donor line (Coriell Institute for Medical Research, GM25256) at passage 37 were electroporated using the 4D-Nucleofector System (Lonza, AAF-1003X, V4XP-3024) with a preassembled complex of 10µg of S.p. Cas9 Nuclease (IDT, 1081058), and 8µg of synthetic sgRNA (Synthego, CRISPRevolution sgRNA EZ Kit,) and 100 pmol of single-stranded oligodeoxynucleotides (ssODNs) (Life Technologies). Cells were replated on Geltrex (Thermo Fisher, A1413301)-coated plates and incubated in mTeSR Plus (Stemcell Technologies, 100- 0276). After five days, single cells were dissociated with Accutase (Innovative Cell Technologies, AT104) sorted into 96-well plates using a WOLF benchtop microfluidic cell sorter (Nanocellect) in mTeSR Plus and CloneR (Stemcell Technologies, 05888). After thirteen days, the genomic DNA was extracted from single clones using QuickExtract solution (Lucigen, QE090500) and HDR was verified by locus specific PCR using DreamTaq DNA Polymerase (Thermo Fisher, EP0702) followed by Sanger sequencing. To generate the BHLHE40 and BHLHE41 double homozygous knockout iPSC line, the cells were electroporated with CRISPR/Cas9 reagents for BHLHE41, and after five days were electroporated again with CRISPR/Cas9 reagents for BHLHE40. One clone per genotype was randomly selected for transcriptional and functional studies. Sequences of guide RNAs, single-stranded oligodeoxynucleotides (ssODNs), and PCR primers used for CRISPR/Cas9-mediated HDR can be found in Supplementary Table 6.

### Genomic integrity of human iPSC lines

The WTC-11 donor human iPSC line (Coriell Institute for Medical Research, GM25256) was tested for genomic integrity at passage 36 using SNP-array technology (Global Diversity Array v1.0 BeadChip, Illumina). No detection of CNV larger than 1.5Mb or AOH larger than 3Mb were detected on somatic chromosomes. The typical WTC-11 deletion of 2.9Mb on Yp11.2 was detected. This deletion is known to be present in the donor, from whom the cell line was derived [95].

### Knockdown of BHLHE40 and/or BHLHE41 by siRNA treatment in human THP-1 macrophages

Human BHLHE40 siRNA, pool of 4 (L-010318-00), human BHLHE41 siRNA, pool of 4 (L- 010043-00), and non-targeting pool (D-001810-10) were purchased from Horizon Discovery Biosciences.

THP-1 monocytes were seeded in 6-well plates at 10^6^ cells per well in RPMI medium supplemented with 10% FBS, 1x Penicillin Streptomycin and 10mM HEPES and 25ng/ml of phorbol-myristate-acetate (PMA) to differentiate monocytes to macrophages (MACs). After 3 days, PMA was removed and replaced with serum-free media (10% FBS was replaced with 1% BSA). Differentiated macrophages were transfected using Lipofectamine RNAiMAX (Thermo Fisher Scientific, LMRNA015) with 20µM *BHLHE40* siRNA (L-010318), 20µM *BHLHE41* siRNA (L-010043), 10µM *BHLHE40* plus 10µM *BHLHE41*, and 20µM of non-targeting control pool (D-001810-10) to generate single (40KD and 41KD) and double (DKD) knock down and scrambled (SCR) groups, respectively. Changes in BHLHE40 and BHLHE41 expression were confirmed at the mRNA level using RT-qPCR and at the protein level using western blotting.

### Generation of human iPSC-derived microglia (iMGLs)

Human induced pluripotent stem cells (iPSCs) were maintained on Matrigel (BD Biosciences) in complete mTeSR Plus (STEMCELL Technologies, 100-0276). iPSCs were passaged every 5-6 days using ReLeSR dissociation reagent (STEMCELL Technologies, 05872) and used for hematopoietic stem cell differentiation with STEMdiff Hematopoietic kit (STEMCELL Technologies, 05310) followed by differentiation to induced microglial-like cells (iMGLs) using a previously published protocol [53]. iPSC lines were confirmed to have a normal karyotype (KaryoStat assay, Thermo Fisher Scientific). iMGLs were maintained and fed with a microglial medium supplemented with three factors (100ng/ml IL34, 50ng/ml TGFβ, 25ng/ml MCSF) for 25 days. On day 25, iMGLs were additionally supplemented with two factors (CX3CL1 and CD200, 100ng/ml each) for an additional three days. Mature iMGLs (day 28) were used for bulk RNA-seq analyses and functional assays.

### Lipid droplet assay

Lipid droplet (LD) quantification was performed using FACS. Cells were collected and stained with 3.7µM BODIPY for 30min at room temperature (RT), protected from light. Single cell data were acquired using Attune flow cytometer (Thermo Fisher Scientific) and analyzed using FCS Express 7 (De Novo Software). Gates were set up based on fluorescence minus one (FMO) controls. LXR agonist TO901317 was purchased from (Sigma-Aldrich, T2320), dissolved in DMSO to 0.01M and used at 10µM final concentration. LXR antagonist GSK2033 was purchased from (Sigma-Aldrich, SML1617), dissolved in DMSO to 5mM and used at 2µM final concentration.

### Cholesterol efflux assay

Cholesterol efflux was performed using Cholesterol Efflux Fluorometric Assay kit (Biovision, K582-100) following manufacturer’s instructions. For this assay, cells were seeded in a 96-well plate at 33,000 cells/well (MACs) or 40,000 cells/well (iMGLs). Briefly, 24h after transfection (MACs) or at day 28 (iMGLs) cells were labeled with Labeling Reagent for 1h at 37°C followed by loading cells with Equilibration Buffer. After overnight incubation, media containing Equilibration Buffer was aspirated and replaced with media containing cholesterol acceptor APOA-1 (10µg/ml) for 4h at 37°C (MACs) or human HDL (25µg/ml) for 4h at 37°C (iMGLs). At the end of incubation, supernatants were transferred to flat bottom clear 96-well white polystyrene microplates (Greiner Bio-one, 655095). Adherent cell monolayers were lysed with Cell Lysis Buffer and incubated for 30min at RT with gentle agitation followed by pipetting up and down to disintegrate cells. Cell lysates were transferred into flat bottom clear 96-well white polystyrene microplates. Fluorescent intensity (Ex/Em=485/523nm) of supernatants and cell lysates was measured using Varioskan LUX multimode microplate reader (Thermo Fisher Scientific, VL0000D0). Percentage of cholesterol efflux was quantified as follows: % cholesterol efflux = Fluorescence intensity of supernatant / fluorescence intensity of supernatant plus fluorescence intensity of cell lysate x 100. We performed a total of 5-6 independent experiments with 2-3 technical replicates per experiment that were averaged within each experiment.

### Lysosomal assays

At day 28 iMGLs cultures were stained with 100nM LysoTracker-Red (Thermo Fisher Scientific, L7525) for 30min at 37°C followed by 1µM LysoSensor-Green (Thermo Fisher Scientific, L7535) staining for 1min at 37°C. To characterize hydrolytic capacity of lysosomes, cells were stained with 1µg/ml DQ Red BSA (Thermo Fisher Scientific, D12051) for 1h at 37°C. Cells were also stained with LIVE/DEAD Fixable Violet Dead Cell Stain (Life Technologies, L34973) to exclude dead cells. After collecting the cells, single cell data were acquired using Attune flow cytometer (Thermo Fisher Scientific) and analyzed using FCS Express 7 (De Novo Software). Gates were set up based on fluorescence minus one (FMO) controls. In addition, we quantified DQ-BSA red fluorescent signal over time using the Incucyte S3 live imaging system. Cells were plated in 96-well plates (40,000 cells/well) and treated with 1µg/ml of DQ Red BSA. Images were acquired every 30min over 5h at 37°C. Total integrated density was calculated as mean red fluorescent intensity multiplied by surface area of masked object (i.e. cell), [RCU x µm^2^].

### Quantification of secreted APOE by ELISA

Secreted APOE was measured using human APOE ELISA kit (Thermo Fisher Scientific, EHAPOE) following manufacturer’s instructions. Briefly, non diluted culture supernatants (100µl) were added to wells coated with human APOE antibody and incubated overnight at 4°C with gentle agitation followed by incubation with solution containing biotin-conjugate (1h at RT) and then solution containing streptavidin-HRP (1h at RT). Reaction was developed using TMB substrate incubated for 30min at RT followed by adding stop solution. Absorbance was read at 450nm using Varioskan LUX multimode microplate reader (Thermo Fisher Scientific, VL0000D0). Sample concentrations were quantified based on the standard curve and normalized to total protein obtained through BCA protein assay kit (Thermo Fisher Scientific, 23225) We performed 5-6 independent experiments with 2-3 technical replicates per experiment that were averaged within each experiment.

### Western blotting

Cells were lysed in NE-PER kit to extract nuclear and cytoplasmic fractions separately (Thermo Fisher Scientific, 78833) supplemented with Protease/Phosphatase Inhibitor Cocktail (Cell Signaling, 5872) following manufacturer’s instructions. Protein concentration was measured using BCA kit (Thermo Fisher Scientific, 23225) and equal quantities were used to prepare samples for western blotting. Samples were resolved by electrophoresis with Bolt 4–12% Bis-Tris Plus Gels (Invitrogen) in Bolt MES SDS running buffer (Invitrogen, B0002) and transferred using iBlot 2 nitrocellulose transfer stacks (Invitrogen). Membranes were blocked for 1h and probed with antibodies: BHLHE40 1:500 (Thermo Fisher Scientific, PA5-83044), BHLHE41 1:500 (Biorbyt, orb224120), APOE 1:1000 (Millipore, AB947), ABCA1 1:1000 (Abcam, 18180), ACTIN 1:10000 (Sigma-Aldrich, A5441) in 5% non-fat dry milk in PBS / 0.1% Tween-20 buffer overnight at 4°C. Secondary antibody staining 1:10000 was applied for 1h at RT, visualized using WesternBright ECL HRP Substrate Kit (Advansta, K-12045), and measured using UVP ChemiDoc-ItTS2 Imager (UVP). Images were analyzed using ImageJ (NIH).

### Quantification of gene expression by RT-qPCR

iMGLs (40, 41KO, KO, DKO and WT as control) were collected on day 28. MACs (40KD, 41KD, DKD and SCR as control) were collected 48h after siRNA transfection. Cell pellets were used for mRNA extraction using the RNeasy Plus Mini kit (Qiagen, 74136) following manufacturer’s instructions. mRNA quantity was measured using Nanodrop 8000 (Thermo Fisher Scientific) and reverse transcription reaction was performed with 1000ng of RNA using High-Capacity RNA-to-cDNA kit (Thermo Fisher #4387406). 10ng cDNA was used in the qPCR reaction with Power SYBR Green Master Mix (Applied Biosystems, 4368706) run using QuantStudio 7 Flex Real-Time PCR System (Thermo Fisher Scientific). Primers were designed using the IDT software and are listed in Supplementary Table 6. Ct values were averaged from two technical replicates for each gene, *GAPDH* Ct values were used for normalization. Gene expression levels were quantified using the 2-^ddCt^ method relative to control. We performed a total of 6-7 independent experiments with 3 technical replicates (i.e. separate wells) per experiment. Technical replicates were averaged within each experiment.

### Quantification of secreted cytokines by antibody array

Conditioned media (1ml) was collected, cleared of cell debris with 10min centrifugation at 4000rpm and used for the Proteome Profiler Human Cytokine Array Kit, Panel A (R&D Systems, ARY005B) following manufacturers’ instructions.

### Statistical analysis

Data were analyzed and visualized in GraphPad Prism 9 (GraphPad Software). In each analysis, three to six independent experiments were performed. The researcher was not blinded to siRNA treatment or genotype. Differences of means between groups were tested with one-way repeated measures ANOVA followed by Dunnett’s post-hoc tests. All data are represented as group mean ± standard error of the mean (SEM). Detailed statistics are shown in Supplementary File 1.

### Acute isolation of microglia from Bhlhe40/41 double knockout mice

Female Bhlhe40^−/−^Bhlhe41^−/−^ (double knock-out, DKO) or wildtype (C57BL/6J or Rosa26^Stop-YFP^) mice were transcardially perfused with 10ml cold PBS prior to brain collection. Olfactory bulb and cerebellum were discarded, followed by enzymatic digestion of the remaining brain in IMDM medium (Cytiva, SH30259.02) supplemented with 1mg/ml Collagenase type IV (Worthington, LS004186) and 33.3U/ml DNase I (Roche, 11284932001) at 37°C for 45min with occasional mixing and dissociation by pipetting. Subsequently, cells were kept on ice or at 4°C throughout further processing. Enzymatic digestion was followed by filtering through a 70µm cell strainer, washing with ice-cold 2% FCS/PBS and spinning at 300xg for 10min. To remove myelin, the pellet was resuspended in 38% percoll/PBS (Cytiva, GE17-0891-02) and spun at 800xg for 15min (no break). Cells were washed in ice-cold 2% FCS/PBS, followed by staining with the following antibodies in ice-cold 2% FCS/PBS: CD38-PE (Miltenyi, 130-103-008), Ly6G- APC (Miltenyi, 130-120-803), Ly6C-BV510 (Biolegend, 127633), CD11b-FITC (Biolegend, 101206), MHCII-BV421 (Biolegend, 107632), CD45.2-PE-Cy7 (Biolegend, 109830), and FcR block (in-house, clone 2.4G2). Cells were washed in PBS, followed by staining with fixable viability dye eFluor 780 (eBioscience, 65-0865-14). Microglia were double sorted on a BD FACSAria Fusion using the following gating strategy: viability dye^-^ Ly6C^-^Ly6G^-^MHCII^-^CD38^-^CD45.2^+^CD11b^+^. Immediately after completion of the sort, cells were pelleted, washed with ice-cold PBS, and cells were disrupted by adding RLT plus buffer with β-mercaptoethanol following manufacturer’s instructions (Qiagen, 1053393), followed by storage at -80°C until further processing.

### RNA-seq analysis

RNA from human iPSC-derived microglia (iMGLs), human THP-1 macrophages (MACs) and acutely isolated mouse microglia was extracted using the RNeasy Plus Mini kit (Qiagen, 74136) following manufacturer’s instructions. RNA was submitted to GENEWIZ (New Jersey, NJ, USA) for QC, library preparation, and next-generation sequencing. Samples passed quality control with Qubit and BioAnalyzer showing RIN > 7.8. RIN for mouse samples were not assessed due to low RNA abundance. Libraries were prepared using TruSeq RNA Sample Prep Kit v2 and paired-end sequenced using HiSeq2500 at a read length of 150bp to obtain 20-30M mapped fragments per sample. Sequenced reads were assessed for quality (FastQC v0.11.8), trimmed for adapter contamination (Cutadapt v2.6), and aligned to the mouse genome mm10 or human genome hg38 (STAR v2.5.3a). Differential gene expression analysis (DGEA) was performed using a linear mixed model implemented in dream (differential expression for repeated measures, variancePartition R package v1.23.1 and R v3.5.3 [96]). Genes with FDR Adj.P-value < 0.05 were considered differentially expressed (DEGs). To identify pathways enriched in human iMGLs lacking BHLHE40 and/or BHLHE41, THP-1 macrophages with reduced levels of *BHLHE40* and/or *BHLHE41,* and mouse microglia lacking *Bhlhe40/41*, we used Gene Set Enrichment Analysis (GSEA) [97]. Briefly, ranked lists were generated from differential gene expression analyses by ordering genes according to the signed test statistic. This metric takes into account both the fold change across conditions as well as the standard error of the fold change. Ranked lists were analyzed using the “GSEA Preranked” module using default GSEA settings including 1000 permutations. Our preranked lists were tested for enrichment against genesets from the Molecular Signatures Database (MSigDB v7.5.1, Broad Institute, C5.all.v2022.1.Hs.symbols.gmt). Enrichment scores were normalized by geneset size to generate normalized enrichment scores (NES) according to the standard protocol [97] and these normalized enrichment scores were used to determine significance of enrichment [10]. DGEA and GSEA results are shown in Supplementary Tables (see Supplementary Table 3 for human iPSC-derived microglia, Supplementary Table 4 for human THP-1 macrophages, and Supplementary Table 5 for acutely isolated mouse microglia).

### Rank-rank hypergeometric overlap (RRHO) and Ingenuity pathway (IPA) analyses

Transcriptional signatures from human lipid-associated macrophages (Jaitin *et al.* [54], Dataset S6, FDR Adj.P-value < 0.05) and from BHLHE40-KO, BHLHE41-KO, and BHLHE40/41-DKO human iPSC-derived microglia (iMGLs), BHLHE40-KD, BHLHE41- KD, BHLHE40/41-DKD human THP-1 macrophages as well as from mouse disease-associated microglia (Keren-Shaul *et al.* [10], Table S3, FDR Adj.P-value < 0.05) and mouse Bhlhe40/41-DKO were compared pairwise using the RRHO2 R package [55,56,98]. The recommended *-log10(P-value) * sign(log2FC)* metric was used to generate ranked lists of genes for each transcriptional signature. RRHO2 was then used to visualize both concordant and discordant gene expression changes across each pair of signatures as rank-rank hypergeometric overlap (RRHO) heatmaps. The color temperature of each pixel in an RRHO heatmap represents the negative log10- transformed hypergeometric overlap test P-value of subsections of the two ranked gene lists, adjusted for multiple testing using the Benjamini-Hochberg correction method. Heatmaps generated using RRHO2 have top-right (both decreasing) and bottom-left (both increasing) quadrants, representing concordant gene expression changes, while the top-left and bottom-right quadrants represent discordant gene expression changes. The default step size and P-value representation method (*hyper*) were used. Gene lists that provide the most significant overlap (expression changes going down or up in compared signatures) were retrieved using overlap_uu or overlap_dd options. Pathway (biological functions and diseases) enrichment and activity analyses of overlapping genes were performed using Ingenuity Pathway Analysis (Qiagen Inc., https://digitalinsights.qiagen.com/IPA) using Z statistics of differential gene expression retrieved from bulk RNA-seq analyses (Supplementary Table 2)

## Availability of data and materials

All aligned read counts and FASTQ files for 40KO, 41KO, and DKO human iPSC-derived microglia (iMGLs), 40KD, 41KD, and DKD human THP-1 macrophages (MACs), as well as microglia acutely isolated from the brain of *Bhlhe40* and *Bhlhe41* double knockout (DKO) mice will be deposited to the Gene Expression Omnibus once the manuscript is accepted for publication. iPSC lines lacking BHLHE40 and/or BHLHE41 will be available upon request.

## Abbreviations

AD: Alzheimeir’s disease
LAM: Lipid-associated macrophages
GRN: Gene regulatory network
qPCR: Quantitative polymerase chain reaction
RNAseq: Ribonucleic acid sequencing
LD: Linkage disequilibrium
IPA: Ingenuity pathway analysis
iPSC: Induced pluripotent stem cell
iMGLs: iPSC-derived microglia
LDs: Lipid droplets
ABCA1: ATP binding cassette subfamily A member 1
ABCG1: ATP binding cassette subfamily G member 1
APOE: Apolipoprotein E
CLU: Clusterin
TREM2: Triggering receptor expressed on myeloid cells 2
SPP1: Secreted phosphoprotein 1
LPL: Lipoprotein lipase
GPNMB: Glycoprotein NMB
CTSZ: Cathepsin Z
CTSB: Cathepsin B
NR1H3: Nuclear Receptor Subfamily 1 Group H Member 3
LXRα: Liver X Receptor alpha
RXR: Retinoid X Receptor
BHLHE40: Basic Helix-Loop-Helix family member E40
BHLHE41: Basic Helix-Loop-Helix family member E41
scRNAseq: single cell Ribonucleic acid sequencing
snRNAseq: single nuclei Ribonucleic acid sequencing
WB: Western blotting
ELISA: Enzyme Linked Immunosorbent Assay
MI: Mutual interactions

## Authors’ contributions

Conceptualization, study design and methodology: G.N., A.P-D., A.M.G., E.M.; data collection, analysis and visualization: G.N., A.P-D.; sample/dataset generation: R.T. (iMGLs), J.D.,T.K. (mouse microglia), C.G. (human atherosclerotic macrophage dataset); strategy for CRISPR/Cas9 genome editing: S.M.; RNA-seq data processing: Y.L.; writing of original draft, G.N., A.P-D.; writing, review, revising, A.M.G., E.M., T.K.. All authors read and approved the final manuscript.

---

**Supplementary Figure 1.**
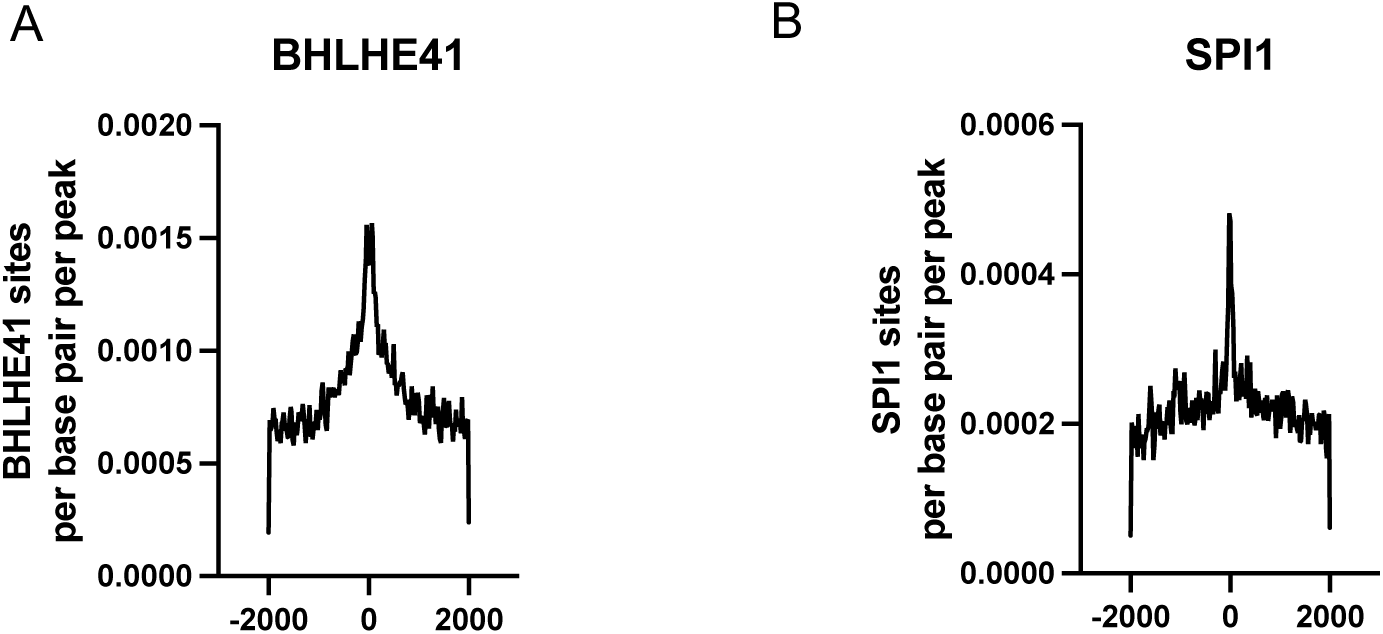
BHLHE41 and SPI1/PU.1 regulons are most highly enriched in direct targets. Frequency of **A)** BHLHE41 and **B)** SPI1/PU.1 motifs are plotted against the distance from the microglial ATAC-seq peaks[31] in the promoters of genes in their regulons.

**Supplementary Figure 2.**
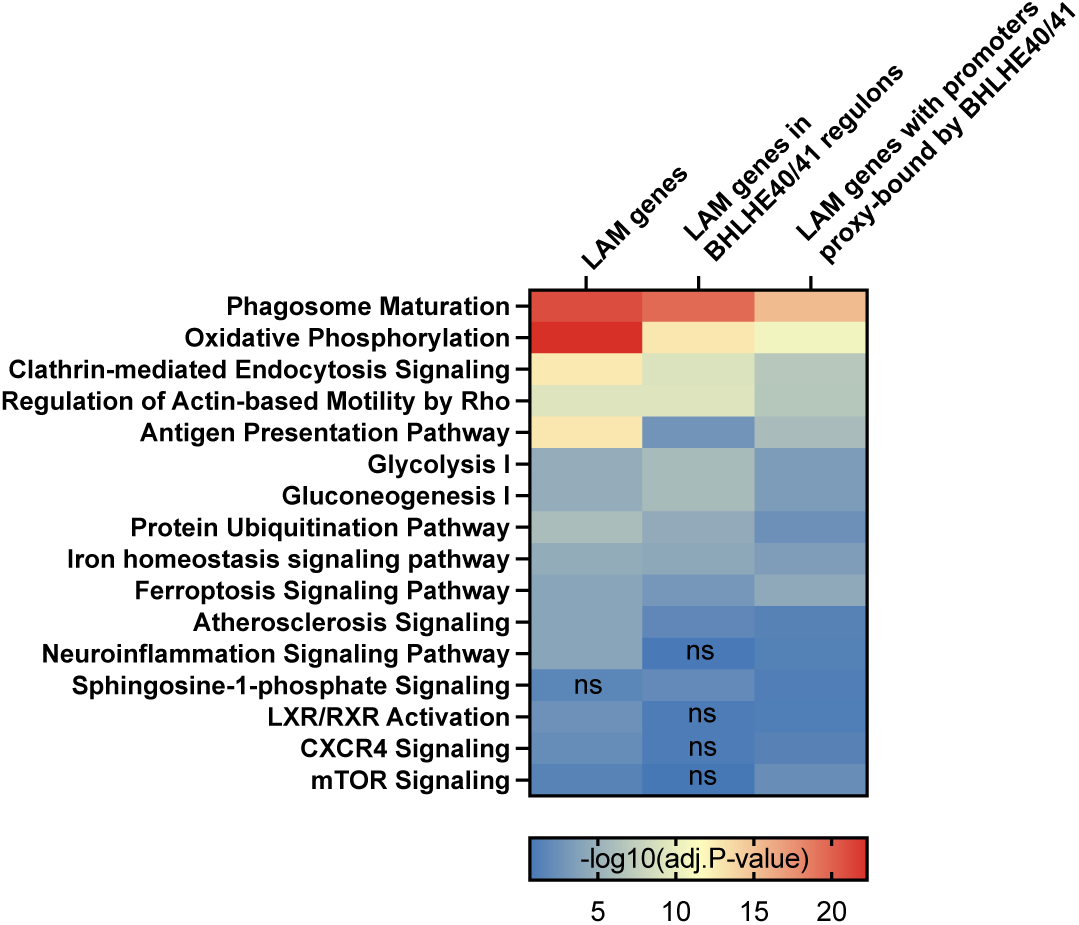
Pathways shared between lipid-associated macrophage (LAM) genes with promoter proxy-bound by BHLHE40/41 and LAM genes in BHLHE40/41 regulons. Canonical pathways (IPA) shared between human LAM genes (Jaitin *et al*. [3], Dataset S6, FDR Adj.P-value < 0.05), LAM genes proxy-bound by BHLHE40/41, and LAM genes from BHLHE40/41 regulons. Adj.P-value represents Bonferroni-Holm correction. ns – non significant (Adj.P-value > 0.05).

**Supplementary Figure 3.**
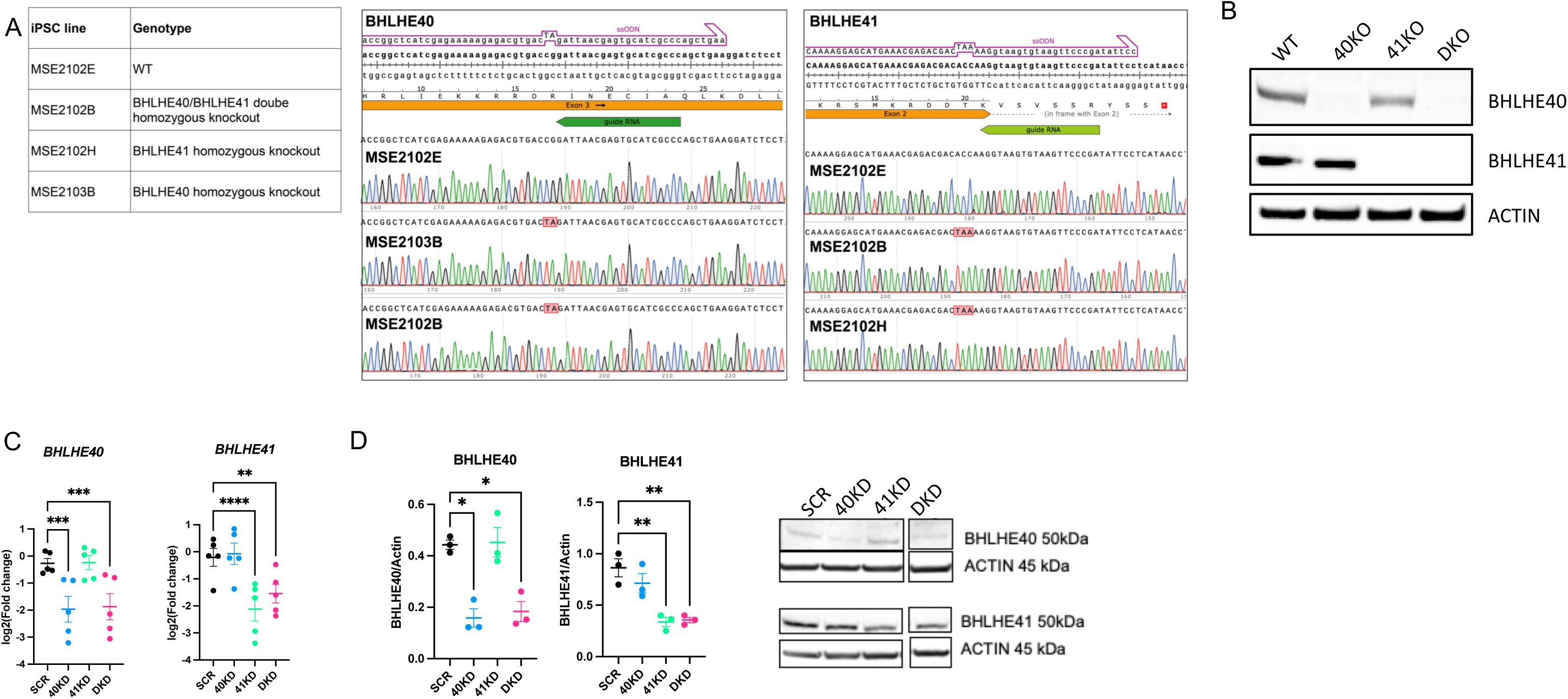
Validation of knockout (KO) efficiency in human iPSC-derived microglia (iMGLs) with genetic inactivation of BHLHE40 and/or BHLHE41 and knockdown (KD) efficiency in human THP-1 macrophages (MACs) treated with BHLHE40 and/or BHLHE41 siRNAs. **A)** CRISPR/Cas9-mediated genome editing strategy to obtain homozygous BHLHE40 and/or BHLHE41 knockout human iPSC lines (Methods). **B)** Western blot confirming loss of BHLHE40 in 40KO and DKO iMGLs and loss of BHLHE41 in 41KO and DKO iMGLs. **C)** Expression of BHLHE40 and BHLHE41 measured by RT-qPCR, N=5/group. log2(fold change) (log2FC) is calculated with SCR MACs as reference. **D)** BHLHE40 and BHLHE41 normalized to Actin measured by western blot, N=3/group. Differences of means between groups were tested using one-way ANOVA with repeated measures followed by Dunnett’s post-hoc test. * P-value < 0.05, ** P-value < 0.01, *** P-value < 0.001, **** P-value < 0.0001. Data plotted as mean ± SEM. Detailed statistics are shown in Supplementary File 1. 40KO = BHLHE40 KO iMGLs, 41KO = BHLHE41 KO iMGLs, DKO = BHLHE40 and BHLHE41 double KO iMGLs, WT = iMGLs derived from the parental iPSC line. 40KD = MACs treated with BHLHE40 siRNA, 41KD = MACs treated with BHLHE41 siRNA, DKD = MACs treated with BHLHE40 and BHLHE41 siRNA, SCR = MACs treated with scrambled siRNA.

**Supplementary Figure 4.**
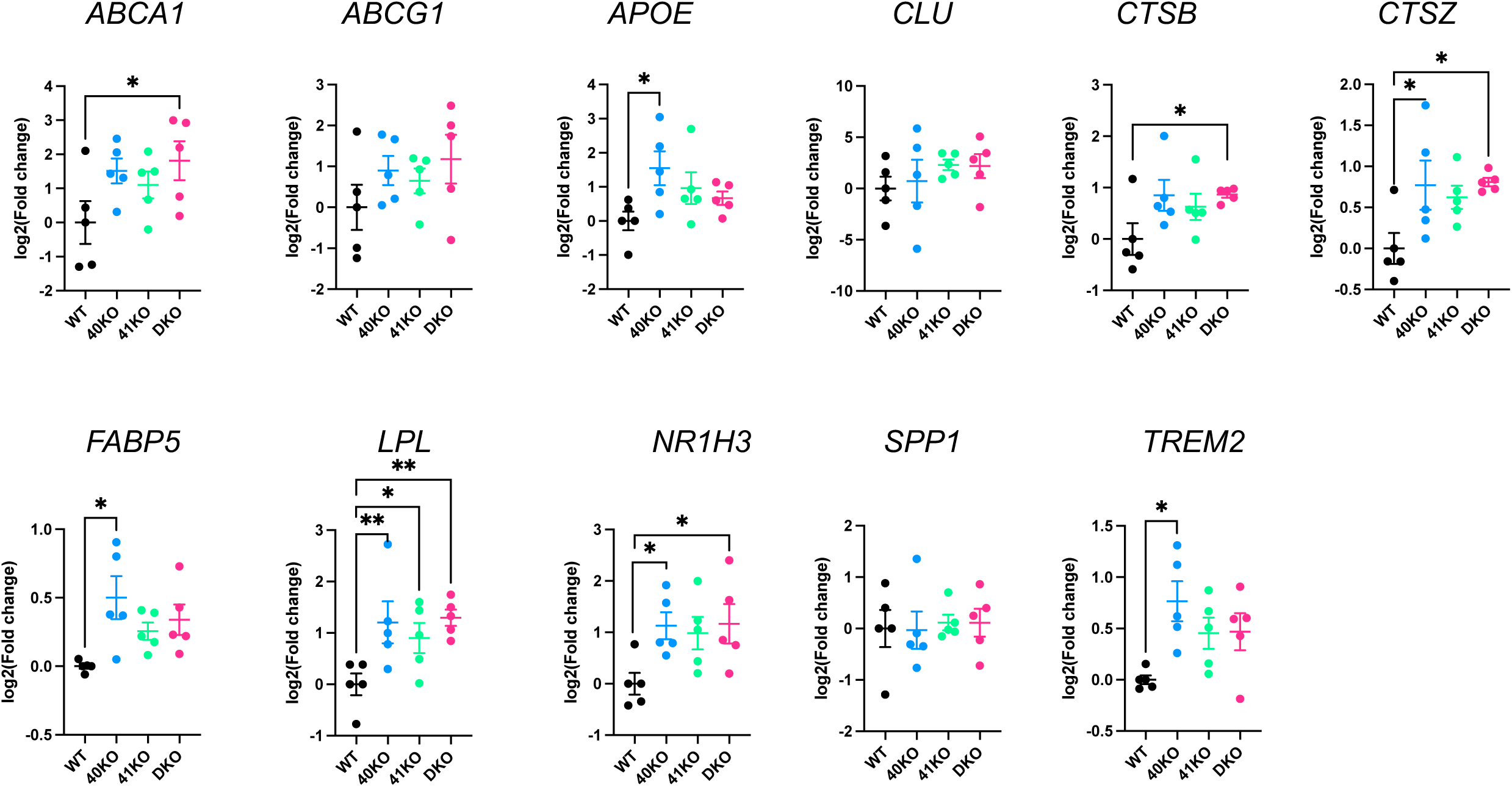
Expression of lipid and lysosomal clearance genes in human iPSC-derived microglia (iMGLs) lacking *BHLHE40* and/or *BHLHE41*. Expression of lipid and lysosomal clearance genes measured by RT-qPCR, N=5/group. log2(fold change) (log2FC) is calculated with WT iMGLs as reference. Differences of means between groups were tested using one-way ANOVA with repeated measures followed by Dunnett’s post-hoc test. * P-value < 0.05, ** P-value < 0.01. Data plotted as mean ± SEM. Detailed statistics are shown in Supplementary File 1. 40KO = BHLHE40 KO iMGLs, 41KO = BHLHE41 KO iMGLs, DKO = BHLHE40 and BHLHE41 double KO iMGLs, WT = iMGLs derived from the parental iPSC line.

**Supplementary Figure 5.**
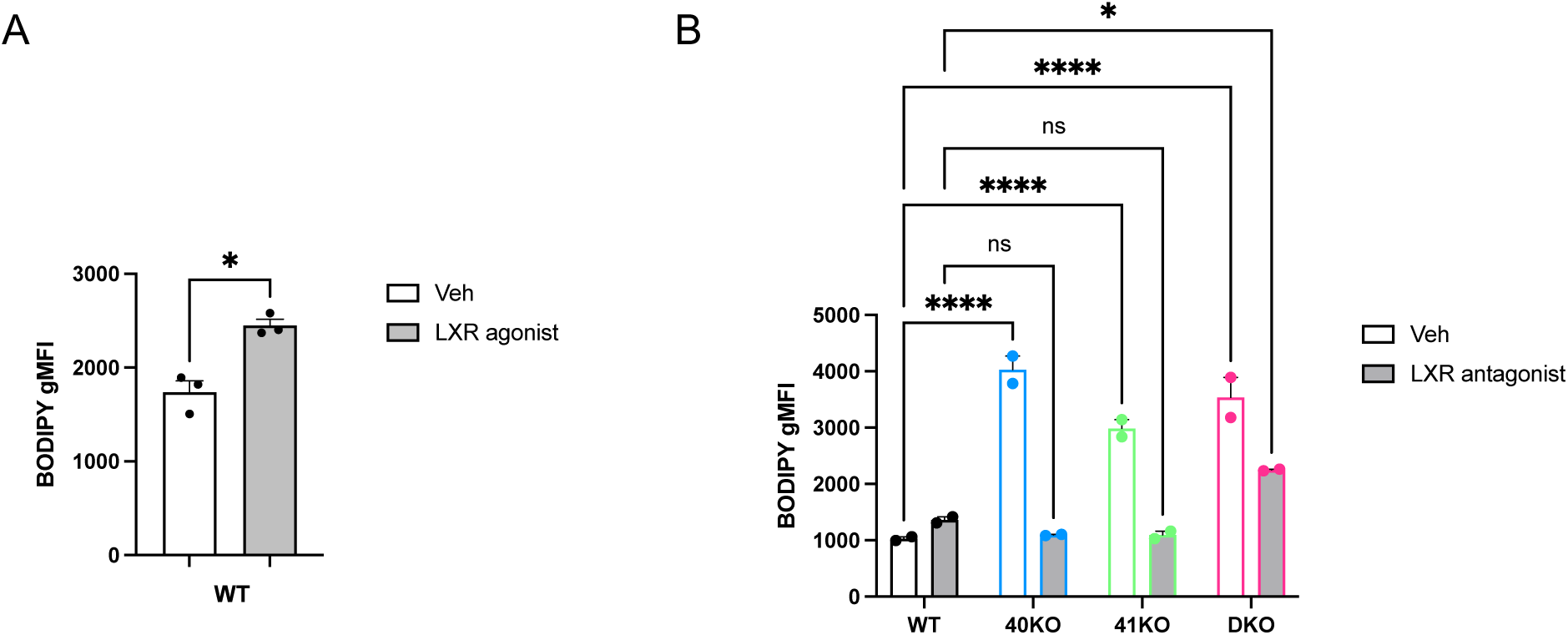
LXR-dependent accumulation of lipid droplets (LDs) in human iPSC-derived microglia (iMGLs) lacking BHLHE40 and/or BHLHE41. **A)** LD content (BODIPY gMFI) measured by flow cytometry in BODIPY-positive WT iMGLs upon treatment with an LXR agonist (TO901317, 10uM, 48h) compared to vehicle control (Veh, DMSO), N=3/group. Differences of means between groups were tested using the paired t.test. * P-value < 0.05. B) LDs content (BODIPY gMFI) measured by flow cytometry in BODIPY-positive WT, 40KO, 41KO, and DKO iMGLs treated with an LXR antagonist (GSK2033, 2uM, 24h) compared to vehicle control (Veh, DMSO), N=2/group. Differences of means between groups were tested using two-way ANOVA followed by Dunnett’s post-hoc tests. Data plotted as mean ± SEM. Detailed statistics are shown in Supplementary File 1. 40KO = BHLHE40 KO iMGLs, 41KO = BHLHE41 KO iMGLs, DKO = BHLHE40 and BHLHE41 double KO iMGLs, WT = iMGLs derived from the parental iPSC line.

**Supplementary Figure 6.**
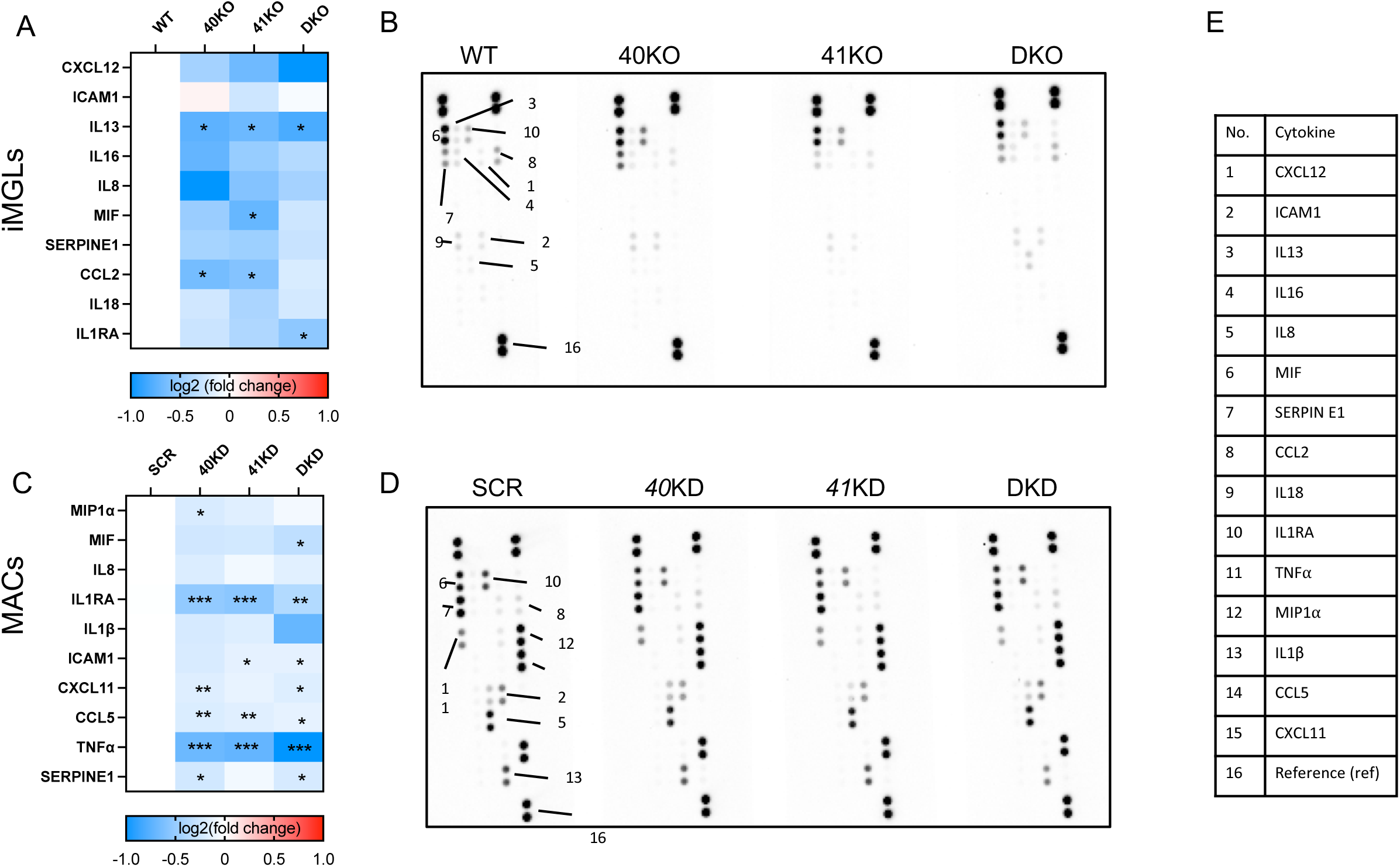
Reduce secretion of proinflammatory cytokines associated with complete loss (KO) or reduced levels (KD) of BHLHE40 and/or BHLHE41 in human iPSC-derived microglia (iMGLs) and THP-1 macrophages (MACs). Quantifications and dot blots of cytokines secreted in conditioned media from (A and B) human iPSC-derived microglia (iMGLs) with genetic inactivation of BHLHE40 and/or BHLHE41 or (C and D) human THP-1 macrophages (MACs) treated with BHLHE40 and/or BHLHE41 siRNAs measured using the Proteome Profiler Human Cytokine Array Kit (R&D Systems). A and C) Levels of cytokines secreted in conditioned media quantified as target density (i.e. mean target spot intensity multiplied by target spot area) normalized to reference density (i.e. mean reference spot intensity multiplied by reference spot area). log2(fold change) is calculated with (A) WT iMGLs or (C) SCR MACs as reference, N=3/group. B and D) Representative dot blots. E) Table of measured cytokines with corresponding dot identification number. Differences of means between groups were tested using one-way ANOVA with repeated measures followed by Dunnett’s post-hoc test. *P-value<0.05, **P-value<0.01, ***P<0.001, ****P< 0.0001. Detailed statistics are shown in Supplementary File 1. 40KO = BHLHE40 KO iMGLs, 41KO = BHLHE41 KO iMGLs, DKO = BHLHE40 and BHLHE41 double KO iMGLs, WT = iMGLs derived from the parental iPSC line. 40KD = MACs treated with BHLHE40 siRNA, 41KD = MACs treated with BHLHE41 siRNA, DKD = MACs treated with BHLHE40 and BHLHE41 siRNA, SCR = MACs treated with scrambled siRNA.

**Supplementary Figure 7.**
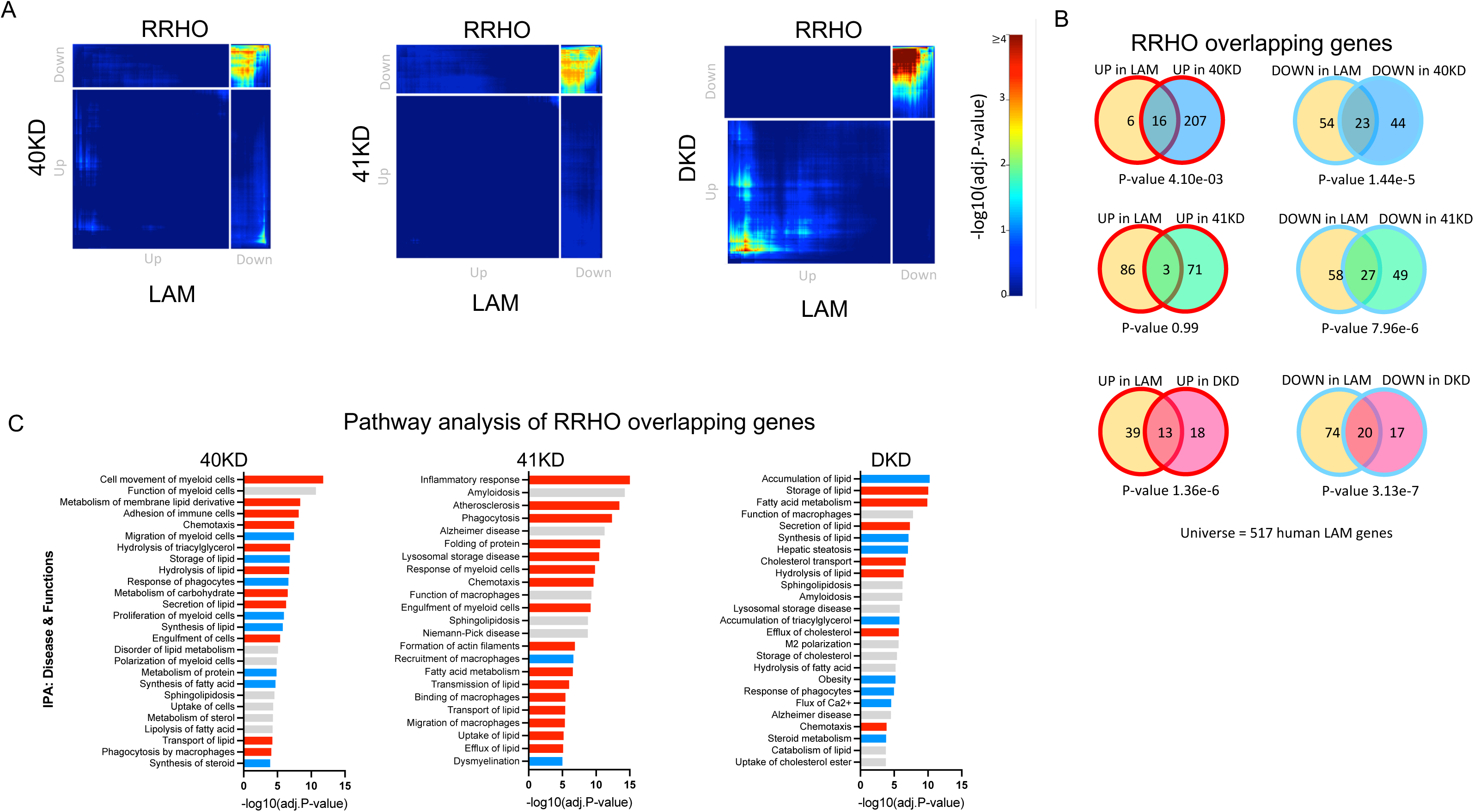
Knockdown of BHLHE40/41 partially recapitulates the LAM transcriptional response in human THP-1 macrophages (MACs). **A)** Rank-rank hypergeometric overlap (RRHO) heatmaps visualizing significant overlaps in gene expression changes between each pair of BHLHE40/41 knockdown (KD) MAC and human LAM (Jaitin et al. [3], Dataset S6, FDR Adj.P-value < 0.05) transcriptional signatures. Adj.P-value in color temperature scale represents Benjamini-Hochberg corrected P-value of hypergeometric overlap test. **B)** Venn diagrams of most significant overlaps between genes up-regulated in both *BHLHE40/41* KD MAC and human LAM transcriptional signatures, corresponding to the warmest pixel in the bottom-left quadrant of the respective heatmap (left) and Venn diagrams of most significant overlaps between genes down-regulated in both BHLHE40/41 KD MAC and human LAM transcriptional signatures, corresponding to the warmest pixel in the upper-right quadrant of the respective heatmap (right). P-values are calculated using the hypergeometric overlap test restricted to the universe of 517 human LAM genes (Jaitin *et al*. [3], Dataset S6, FDR Adj.P-value < 0.05). **C)** Pathways (“diseases and biological functions” category) found by IPA to be significantly enriched for RRHO overlapping genes. Red bars represent pathways predicted by IPA to be more active (i.e., with positive Z-scores), blue bars represent pathways predicted by IPA to be less active (i.e., with negative Z-scores), grey bars represent pathways with non-attributed Z-scores. Adj.P-value on the x-axis represents Bonferroni-Holm corrected P-value of pathway enrichment test. 40KD = MACs treated with BHLHE40 siRNA, 41KD = MACs treated with BHLHE41 siRNA, DKD = MACs treated with BHLHE40 and BHLHE41 siRNA, SCR = MACs treated with scrambled siRNA.

**Supplementary Figure 8.**
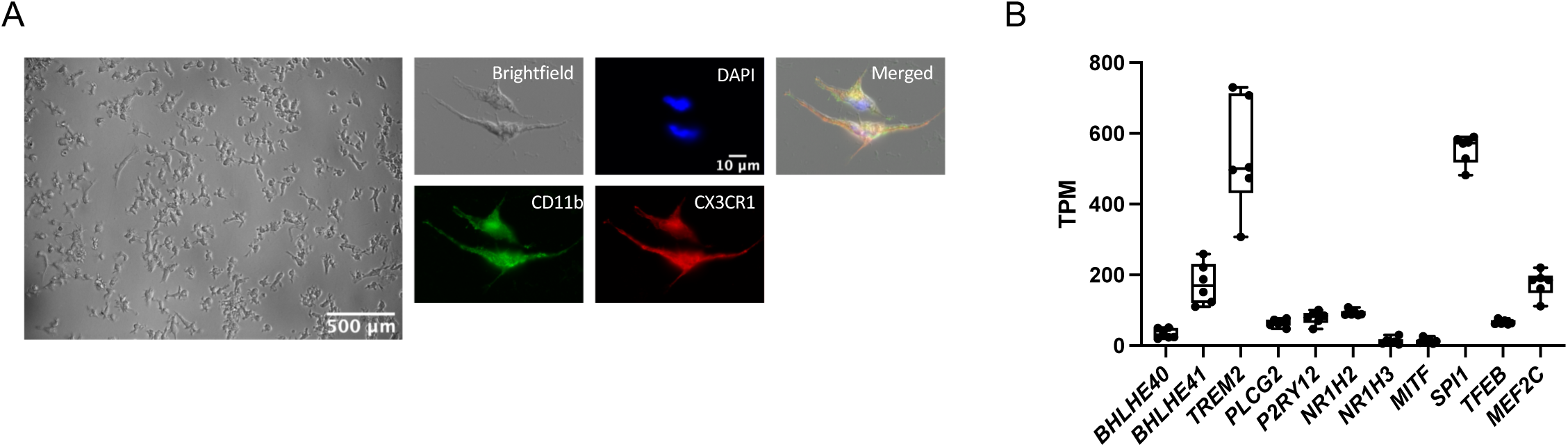
Differentiation of human iPSC lines into microglia-like cells (iMGLs). **A)** Representative bright-field microscopic images and immunofluorescence staining for nuclei (DAPI) and differentiation markers (CD11b and CX3CR1) of a typical culture of mature iMGLs after 25d *in vitro*. **B)** Expression of *BHLHE40* and *BHLHE41* and other microglial genes (*P2RY12*, *PLCG2*, and *TREM2*) and transcription factors (*SPI1*, *MITF*, *MEF2C*, *NR1H2*, *NR1H3*, and *TFEB*) of interest measured by RNA-seq in iMGLs derived from the parental iPSC line, N=5/group. TPM = transcripts per million.

## Figure legends with detailed statistical tests used in Figure 5, Figure 6, Supplementary Figure 3, Supplementary Figure 4, Supplementary Figure 5, Supplementary Figure 6

**Figure 5. Knockout of BHLHE40/41 increases expression of lipid and lysosomal clearance genes, cholesterol efflux and lipid droplet content, lysosomal mass and degradative capacity in human iPSC-derived microglia (iMGLs).**

A. **Expression of lipid and lysosomal clearance genes measured by RT-qPCR.** from N=5 independent iMGL differentiations/group (separate plots and statistical details for each gene in Supplementary Figure 5 and in the legend for Supplementary Figure 5 below). Post-hoc tests * P-value < 0.05, ** P-value < 0.01

B. **Intracellular APOE normalized to Actin measured by western blot.** Differences of means between groups were tested with one-way repeated measures ANOVA F(3,6)=10.66, P-value=0.0081, N=3 independent iMGL differentiations/group, followed by Dunnett’s post-hoc tests: ES_40KO-WT_=1.04 [0.43, 1.68], P-value_40KO- WT_=0.0050; ES_41KO-WT_=0.46 [-0.48, 0.77], P-value_41KO-WT_=0.8204; ES_DKO-WT_=0.98 [- 0.28, 0.97], P-value_DKO-WT_=0.2906.

C. **ABCA1 normalized to Actin measured by western blot.** Differences of means between groups were tested with one-way repeated measures ANOVA F(3,6)=13.11, P-value=0.0048, N=3 independent iMGL differentiations/group, followed by Dunnett’s post-hoc tests: ES_40KO-WT_=0.17 [-0.04, 0.38], P-value_40KO- WT_=0.1068; ES_41KO-WT_=0.42 [0.21, 0.63], P-value_41KO-WT_=0.0021; ES_DKO-WT_=0.14 [- 0.06, 0.35], P-value_DKO-WT_=0.1701.

D. **Secreted APOE normalized to total protein measured by ELISA.** Differences of means between groups were tested with one-way repeated measures ANOVA F(3,9)=6.41, P-value=0.0130, N=5 independent iMGL differentiations/group, followed by Dunnett’s post-hoc tests: ES_40KO-WT_=5.76 [2.03, 9.49], P-value_40KO- WT_=0.0048; ES_41KO-WT_=3.53 [-0.19, 7.23], P-value_41KO-WT_=0.0630; ES_DKO-WT_=2.93 [- 0.79, 6.66], P-value_DKO-WT_=0.1271.

E. **Percentage of cholesterol efflux.** Differences of means between groups were tested with one-way repeated measures ANOVA F(3,12)=24.68, P-value<0.0001, N=6 independent iMGL differentiations/group, followed by Dunnett’s post-hoc tests: ES_40KO-WT_=6.80 [3.76, 9.84], P-value_40KO-WT_=0.002; ES_41KO-WT_=7.80 [4.76, 10.84], P-value_41KO-WT_<0.0001; ES_DKO-WT_=8.80 [5.76, 11.84], P-value_DKO-WT_<0.0001.

A. F. **LD content (BODIPY gMFI) measured by flow cytometry in BODIPY-positive cells** (top). Differences of means between groups were tested with one-way repeated measures ANOVA F(3,15)=7.23, P-value=0.0032, N=6 independent iMGL differentiations/group, followed by Dunnett’s post-hoc tests: ES_40KO-WT_=1928 [783.7, 3072], P-value_40KO-WT_=0.0014; ES_41KO-WT_=1151 [7.441, 2295], P-value_41KO- WT_=0.0484; ES_DKO-WT_=1546 [401.8, 2690], P-value_DKO-WT_=0.0082. Representative flow-cytometry histograms (bottom)

B. G. **Lysosomal mass (LysoTracker gMFI) measured by flow cytometry in LysoTracker-positive cells** (top). Differences of means between groups were tested with one-way repeated measures ANOVA F(3,12)=2.81, P-value=0.0845, N=5 independent iMGL differentiations/group, followed by Dunnett’s post-hoc tests: ES_40KO-WT_=1987 [-1982, 5957], P-value_40KO-WT_=0.4270; ES_41KO-WT_=3125 [- 844.9, 7094], P-value_41KO-WT_=0.1341; ES_DKO-WT_=4080 [109.9, 8049], P-value_DKO- WT_=0.0438. Representative flow-cytometry histograms (bottom)

C. H. **Lysosomal acidification (LysoSensor gMFI) measured by flow cytometry in LysoSensor-positive cells** (top). Differences of means between groups were tested with one-way repeated measures ANOVA F(3,12)=4.41, P-value=0.0261, N=5 independent iMGL differentiations/group, followed by Dunnett’s post-hoc tests: ES_40KO-WT_=1327 [-19426, 22080], P-value_40KO-WT_=0.9963; ES_41KO-WT_=2858 [- 17895, 23611], P-value_41KO-WT_=0.9661; ES_DKO-WT_=24256 [3502, 45009], P- value_DKO-WT_=0.0222. Representative flow-cytometry histograms (bottom)

I. **Lysosomal proteolysis (DQ-BSA gMFI) measured by flow cytometry in DQ-BSA-positive cells** (top). Differences of means between groups were tested with one-way repeated measures ANOVA F(3,12)=9.48, P-value=0.0017, N=5 independent iMGL differentiations/group, followed by Dunnett’s post-hoc tests:

ES40KO-WT=3846 [-961.2, 8653], P-value40KO-WT=0.1265; ES41KO-WT=3125 [5.313, 9620], P-value41KO-WT=0.0497; ESDKO-WT=9485 [4678, 14293], P-valueDKO-_WT_=0.0005. Representative flow-cytometry histograms (bottom).

A. J. **DQ-BSA red fluorescent signal (total integrated density) measured over time using the Incucyte S3 live imaging system.** Differences of means between groups were tested with two-way repeated measures ANOVA F(3.8)=13.93, P-value=0.0015, N=3 independent iMGL differentiation/group, followed by Dunnett’s post-hoc tests: ES_40KO-WT_=37.34 [-106.7, 181.4], P-value_40KO-WT_=0.7997; ES_41KO- WT_=192 [47.91, 336.0], P-value_41KO-WT_=0.0126; ES_DKO-WT_=281.6 [137.5, 425.6], P- value_DKO-WT_=0.0013. Total integrated density was calculated as mean red fluorescent intensity multiplied by surface area of masked object (i.e. cell), [RCU x µm^2^].

Effect sizes (ES) are reported as unstandardized point estimates with 95% confidence intervals in the same unit as depicted on each graph.

**Figure 6. Knockdown of BHLHE40/41 increases expression of lipid and lysosomal clearance genes, cholesterol efflux and lipid droplet content in human THP1 macrophages (MACs).**

A. **Expression of lipid and lysosomal clearance genes measured by RT-qPCR.** from N=7 independent MAC differentiation (transient transfection with siRNA)/group. Post-hoc tests * P-value < 0.05, ** P-value < 0.01

B. **Secreted APOE normalized to total protein measured by ELISA.** Differences of means between groups were tested with one-way repeated measures ANOVA F(3,15)=2.52, P-value=0.0976, N=6 independent MAC differentiations/transient transfection with siRNA per group, followed by Dunnett’s post-hoc tests: ES_40KD- SCR_=2.33 [0.04, 4.62], P-value_40KO-WT_=0.0457; ES_41KD-SCR_=0.66 [-1.62, 2.94], P-value_41KD-SCR_=0.7914; ES_DKD-SCR_=0.8324 [1.45, 3.12], P-value_DKD-SCR_=0.6689.

C. **Percentage of cholesterol efflux.** Differences of means between groups were tested with one-way repeated measures ANOVA F(3,15)=22.82, P-value<0.0001,

N=6 independent MAC differentiations/transient transfection with siRNA per group, followed by Dunnett’s post-hoc tests: ES40KD-SCR=12.45 [8.47, 16.44], P-value40KO- WT<0.0001; ES41KD-SCR=4.83 [0.84, 8.81], P-value41KD-SCR=0.0170; ES_DKD-SCR_=6.906 [2.92, 10.89], P-value_DKD-SCR_=0.0011.

A. D. **LD content (BODIPY gMFI) measured by flow cytometry in BODIPY-positive cells.** Differences of means between groups were tested with one-way repeated measures ANOVA F(3,15)=10.31, P-value=0.0006, N=6 independent MAC differentiations/transient transfection with siRNA per group, followed by Dunnett’s post-hoc tests: ES_40KD-SCR_=2348 [567.0, 4128], P-value_40KO-WT_=0.0097; ES_41KD- SCR_=2890 [1110, 4671], P-value_41KD-SCR_=0.0020; ES_DKD-SCR_=3563 [1782, 5343], P- value_DKD-SCR_=0.0003. Gates were drown based on fluorescence minus one (FMO) controls.

Effect sizes (ES) are reported as unstandardized point estimates with 95% confidence intervals in the same unit as depicted on each graph.

**Supplementary Figure 3. Validation of knockout (KO) efficiency in human iPSC-derived microglia (iMGLs) with genetic inactivation of BHLHE40 and/or BHLHE41 and knockdown (KD) efficiency in human THP-1 macrophages (MACs) treated with BHLHE40 and/or BHLHE41 siRNAs.**

A. CRISPR-Cas9 genome editing strategy to obtain homozygous BHLHE40 and/or BHLHE41 knockout human iPSC lines.

B. Western blot confirming loss of BHLHE40 in 40KO and DKO iMGLs and loss of BHLHE41 in 41KO and DKO iMGLs.

C. Expression of BHLHE40 and BHLHE41 measured by RT-qPCR, N=5/group. log2(fold change) (log2FC) is calculated with SCR MACs as reference. Differences of means between groups were tested with one-way repeated measures followed by Dunnett’s post-hoc tests *BHLHE40:* ANOVA F(3,12)=20.02, P-value<0.0001, N=5 independent MAC differentiations/transient transfection with siRNA per group, followed by Dunnett’s post-hoc tests: ES40KD-SCR=-1.699 [-2.498, -0.8999], P-value40KO-WT=0.0003; ES41KD-SCR=0.0283 [-0.7707, 0.8274], P-value41KD-SCR=0.9993; ESDKD-SCR=-1.606 [-2.405, -0.8073], P-value_DKD-SCR_=0.0005.

*BHLHE41:* ANOVA F(3,12)=25.21, P-value<0.0001, N=5 independent MAC differentiations/transient transfection with siRNA per group, followed by Dunnett’s post-hoc tests: ES40KD-SCR=0.1356 [-0.6250, 0.8963], P-value40KO-WT=0.9318; ES41KD-SCR=1.912 [-2.673, 1.151], P-value41KD-SCR<0.0001; ESDKD-SCR=-1.338 [-2.098, -0.5770,], P-value_DKD-SCR_=0.0014.

A. D. **BHLHE40 and BHLHE41 normalized to Actin measured by western blot** Differences of means between groups were tested with one-way repeated measures followed by Dunnett’s post-hoc tests

BHLHE40: ANOVA F(3,6)=12.51, P-value<0.0054, N=3 independent MAC differentiations/transient transfection with siRNA per group, followed by Dunnett’s post-hoc tests: ES40KD-SCR=-0.2847 [-0.4832, 0.0862], P-value40KO-WT=0.0107; ES41KD-SCR=-0.0091 [-0.1894, 0.2076], P-value41KD-SCR=0.9977; ESDKD-SCR=-0.2599 [-0.4584, -0.0613], P-valueDKD-SCR=0.0162.

BHLHE41: ANOVA F(3,12)=14.62, P-value=0.0036, N=3 independent MAC differentiations/transient transfection with siRNA per group, followed by Dunnett’s post-hoc tests: ES40KD-SCR=-0.1502 [-0.4509, 0.1504], P-value40KO-WT=0.3576; ES41KD-SCR=-0.5257 [-0.8264, -0.2250], P-value41KD-SCR=0.0041; ESDKD-SCR=-0.5075 [-0.8082, -0.2069], P-value_DKD-SCR_=0.0048.

Effect sizes (ES) are reported as unstandardized point estimates with 95% confidence intervals in the same unit as depicted on each graph.

**Supplementary Figure 4. Expression of lipid and lysosomal clearance genes in human iPSC-derived microglia (iMGLs) lacking *BHLHE40* and/or *BHLHE41*. Expression of lipid and lysosomal clearance genes measured by RT-qPCR, N=5/group. log2(fold change) (log2FC) is calculated with WT iMGLs as reference. Differences of means between groups were tested using one-way ANOVA with repeated measures followed by Dunnett’s post-hoc test.**

*ABCA1*: ANOVA F(3,12)=3.17, P-value=0.0640 followed by Dunnett’s post-hoc tests: ES40KO-WT=1.51 [-0.1786, 3.207], P-value40KO-WT=0.0822; ES41KO-WT=1.102 [-0.5912, 2.794], P-value_41KO-WT_=0.2408; ES_DKO-WT_=1.813 [0.1202, 3.506], P-value_DKO-WT_=0.0356.

*ABCG1*: ANOVA F(3,12)=1.45, P-value=0.2783 followed by Dunnett’s post-hoc tests: ES40KO-WT=0.8994 [-0.6879, 2.487], P-value40KO-WT=0.3357; ES41KO-WT=0.6468 [-0.9405, 2.234], P-value_41KO-WT_=0.5775; ES_DKO-WT_=1.178 [-0.4097, 2.765], P-value_DKO-WT_=0.1637.

*APOE*: ANOVA F(3,12)=3.89, P-value=0.0372 followed by Dunnett’s post-hoc tests: ES40KO-WT=1.546 [0.3100, 2.782], P-value40KO-WT=0.0149; ES41KO-WT=0.9622 [-0.2740, 2.198], P-value41KO-WT=0.1395; ESDKO-WT=0.6654 [-0.5708, 1.902], P-value_DKO-WT_=0.3732.

*CLU*: ANOVA F(3,12)=0.7083, P-value=0.5654 followed by Dunnett’s post-hoc tests: ES40KO-WT=0.7216 [-4.371, 5.814], P-value40KO-WT=0.9633; ES41KO-WT=2.306 [-2.786, 7.399], P-value_41KO-WT_=0.5016; ES_DKO-WT_=2.189 [-2.903, 7.282], P-value_DKO-WT_=0.5395.

*CTSB*: ANOVA F(3,12)=2.620, P-value=0.0988 followed by Dunnett’s post-hoc tests: ES40KO-WT=0.8508 [-0.099, 1.801], P-value40KO-WT=0.0818; ES41KO-WT=0.6270 [-0.3231, 1.577], P-value41KO-WT=0.2319; ESDKO-WT=0.8646 [-0.0548, 1.815], P-value_DKO-WT_=0.0764.

*CTSZ*: ANOVA F(3,12)=4.201, P=0.0301 followed by Dunnett’s post-hoc tests: ES40KO- WT=0.7712 [0.0751, 1.467], P-value40KO-WT=0.0298; ES41KO-WT=0.6248 [-0.0713, 1.321], P- value41KO-WT=0.0810; ESDKO-WT=0.8092 [0.1131, 1.505], P-valueDKO-WT=0.0229.

*FABP5*: ANOVA F(3,12)=3.720, P-value=0.0422 followed by Dunnett’s post-hoc tests: ES40KO-WT=0.5012 [0.0895, 0.9129], P-value40KO-WT=0.0175; ES41KO-WT=0.2558 [-0.1559, 0.6675], P-value41KO-WT=0.2712; ESDKO-WT=0.3398 [-0.0719, 0.7515], P-value_DKO-WT_=0.1129.

*LPL*: ANOVA F(3,12)=7.025, P-value=0.0056 followed by Dunnett’s post-hoc tests: ES40KO-WT=1.207 [0.3591, 2.054], P-value40KO-WT=0.0065; ES41KO-WT=0.9020 [0.0544, 1.750], P-value_41KO-WT_=0.0368; ES_DKO-WT_=1.296 [0.4487, 2.144], P-value_DKO-WT_=0.0039.

*NR1H3*: ANOVA F(3,12)=3.438, P-value=0.0520 followed by Dunnett’s post-hoc tests: ES40KO-WT=1.129 [0.001, 2.258], P-value40KO-WT=0.0500; ES41KO-WT=0.9856 [-0.1442, 2.115], P-value_41KO-WT_=0.0910; ES_DKO-WT_=1.166 [0.0364, 2.296], P-value_DKO-WT_=0.0429.

*SPP1*: ANOVA F(3,12)=0.0833, P-value=0.9678 followed by Dunnett’s post-hoc tests: ES40KO-WT=-0.0308 [-1.027, 0.9655], P40KO-WT=0.9996; ES41KO-WT=0.1156 [-0.8807, 1.112], P-value41KO-WT=0.9790; ESDKO-WT=0.1126 [-0.8837, 1.109], P-value_DKO-WT_=0.9805.

*TREM2*: ANOVA F(3,12)=4.113, P-value=0.0320 followed by Dunnett’s post-hoc tests: ES40KO-WT=0.7654 [0.1746, 1.356], P-value40KO-WT=0.0120; ES41KO-WT=0.4536 [-0.1372, 1.044], P-value41KO-WT=0.1462; ESDKO-WT=0.4688 [-0.1220, 1.060], P-value_DKO-WT_=0.1303.

Effect sizes (ES) are reported as unstandardized point estimates with 95% confidence intervals in the same unit as depicted on each graph.

**Supplementary Figure 5. LXR-dependent accumulation of lipid droplets (LDs) in human iPSC-derived microglia (iMGLs) lacking BHLHE40 and/or BHLHE41.**

A. LD content (BODIPY gMFI) measured by flow cytometry in BODIPY-positive WT iMGLs upon treatment with an LXR agonist (TO901317, 10uM, 48h) compared to vehicle control (Veh, DMSO). Differences of means between groups were tested using the paired t.test. t=7.049, df=2, P-value=0.0195, N=3 independent iMGLs differentiations per group ES_LXR_agonist-Veh_=711.3 [277.2, 1145]

B. LD content (BODIPY gMFI) measured by flow cytometry in BODIPY-positive WT, 40KO, 41KO, and DKO iMGLs treated with an LXR antagonist (GSK2033, 2uM, 24h) compared to vehicle control (Veh, DMSO). Differences of means between groups were tested using two-way ANOVA Genotype: F(3,8)=34.17, P<0.0001, Treatment: F(1,8)=152.9, P-value<0.0001, Genotype x Treatment F(3,8) = 40.04, P-value<0.0001, N=2 independent iMGLs differentiation followed by Dunnett’s post-hoc tests Veh: ES_40KO-WT_=2999 [2325, 3672], P-value_40KO-WT_<0.0001; ES_41KO- WT_=1959 [1286, 2632], P-value_41KO-WT_<0.0001; ES_DKO-WT_=2508 [1835, 3181], P- value_DKO-WT_<0.0001; LXR antagonist: ES_40KO-WT_=-268.2 [-941.1, 405.1], P-value_40KO-WT_=0.5531; ES_41KO-WT_=-265.4 [-938.6, 407.8], P-value_41KO-WT_=0.5602; ES_DKO-WT_=887.1 [213.8, 1560], P-value_DKO-WT_=0.0133

Data plotted as mean ± SEM. 40KO = BHLHE40 KO iMGLs, 41KO = BHLHE41 KO iMGLs, DKO = BHLHE40 and BHLHE41 double KO iMGLs, WT = iMGLs derived from the parental iPSC line. Effect sizes (ES) are reported as unstandardized point estimates with 95% confidence intervals in the same unit as depicted on each graph

Supplementary Figure 6. Reduced secretion of proinflammatory cytokines associated with complete loss (KO) or reduced levels (KD) of BHLHE40 and/or BHLHE41 in human iPSC-derived microglia (iMGLs) and THP-1 macrophages (MACs). Quantifications and dot blots of cytokines secreted in conditioned media from (A and B) human iPSC-derived microglia (iMGLs) with genetic inactivation of BHLHE40 and/or BHLHE41 or (C and D) human THP-1 macrophages (MACs) treated with BHLHE40 and/or BHLHE41 siRNAs measured using the Proteome Profiler Human Cytokine Array Kit (R&D Systems).

A. Levels of cytokines secreted in conditioned media quantified as target density (i.e. mean target spot intensity multiplied by target spot area) normalized to reference density (i.e. mean reference spot intensity multiplied by reference spot area). Data are plotted as log2(fold change) which is calculated with WT iMGLs. Differences of means between groups were tested using one-way ANOVA with repeated measures followed by Dunnett’s post-hoc test. N=3/group

CXCL12: ANOVA F(3,6)=2.42, P-value=0.1645, followed by Dunnett’s post-hoc tests: ES40KO-WT=-0.0181 [-0.2717, 0.2353], P-value40KO-WT=0.9916; ES41KO-WT=-0.0817 [-0.3352, 0.1718], P-value_41KO-WT_=0.6498; ES_DKO-WT_=-0.1990 [-0.4525, 0.0545], P- value_DKO-WT_=0.1163.

ICAM1: ANOVA F(3,6)=0.745, P-value=0.5634, followed by Dunnett’s post-hoc tests: ES40KO-WT=0.0361 [-0.2259, 0.2981], P-value40KO-WT=0.9482; ES41KO-WT=-0.0817 [-0.3438, 0.1802], P-value_41KO-WT_=0.6690; ES_DKO-WT_=-0.1496 [-0.2470, 0.2769], P- value_DKO-WT_=0.9957.

IL13: ANOVA F(3,6)=4.534, P-value=0.0550, followed by Dunnett’s post-hoc tests: ES40KO-WT=-0.2847 [-0.5838, -0.00143], P-value40KO-WT=0.0501; ES41KO-WT=-0.2756 [-0.5747, -0.0023], P-value_41KO-WT_=0.0500; ES_DKO-WT_=-0.3076 [-0.6067, -0.0085], P- value_DKO-WT_=0.0448.

IL16: ANOVA F(3,6)=2.354, P-value=0.1712, followed by Dunnett’s post-hoc tests: ES40KO-WT=-0.2548 [-0.5799, 0.0701], P-value40KO-WT=0.1168; ES41KO-WT=-0.2251 [-0.5501, 0.0999], P-value_41KO-WT_=0.1685; ES_DKO-WT_=-0.1566 [-0.4817, 0.1684], P- value_DKO-WT_=0.3817.

IL8: ANOVA F(3,3)=2.410, P-value=0.2445, followed by Dunnett’s post-hoc tests: ES40KO- WT=-0.2953 [-0.7676, 0.1770], P-value40KO-WT=0.1553; ES41KO-WT=-0.1517 [-0.6240, 0.3206], P-value41KO-WT=0.4876; ESDKO-WT=-0.1172 [-0.5895, 0.3550], P-value_DKO-WT_=0.6382.

MIF: ANOVA F(3,6)=2.065, P-value=0.2064, followed by Dunnett’s post-hoc tests: ES40KO-WT=-0.2606 [-0.7211, 0.1999], P-value40KO-WT=0.2776; ES41KO-WT=-0.3457 [-0.8062, -0.0011], P-value_41KO-WT_=0.0491; ES_DKO-WT_=-0.1342 [-0.5947, 0.3263], P- value_DKO-WT_=0.7083.

SERPINE1: ANOVA F(3,6)=1.079, P-value=0.4264, followed by Dunnett’s post-hoc tests: ES40KO-WT=-0.2159 [-0.6675, 0.2357], P-value40KO-WT=0.3872; ES41KO-WT=-0.2367 [-0.6883, 0.2150], P-value_41KO-WT_=0.3263; ES_DKO-WT_=-0.1515 [-0.6031, 0.3001], P- value_DKO-WT_=0.6255.

CCL2: ANOVA F(3,6)=6.951, P-value=0.0223, followed by Dunnett’s post-hoc tests: ES40KO-WT=-0.2371 [-0.4208, -0.0533], P-value40KO-WT=0.0173; ES41KO-WT=-0.2109 [-0.3946, -0.0271], P-value_41KO-WT_=0.0288; ES_DKO-WT_=-0.0887 [-0.2714, 0.0961], P- value_DKO-WT_=0.3887.

IL18: ANOVA F(3,6)=2.551, P-value=0.1129, followed by Dunnett’s post-hoc tests: ES40KO-WT=-0.2271 [-0.4208, 0.0633], P-value40KO-WT=0.1067; ES41KO-WT=-0.1909 [-0.3946, 0.0471], P-value_41KO-WT_=0.1175; ES_DKO-WT_=-0.1887 [-0.2714, 0.00961], P- value_DKO-WT_=0.0763.

IL1RA: ANOVA F(3,6)=2.032, P-value=0.2109, followed by Dunnett’s post-hoc tests: ES40KO-WT=-0.1429 [-0.4931, 0.2073], P-value40KO-WT=0.4957; ES41KO-WT=-0.1878 [-0.5380, 0.1624], P-value_41KO-WT_=0.3114; ES_DKO-WT_=-0.2722 [-0.6224, -0.0077], P- value_DKO-WT_=0.0493.

C. Levels of cytokines secreted in conditioned media quantified as target density (i.e. mean target spot intensity multiplied by target spot area) normalized to reference density (i.e. mean reference spot intensity multiplied by reference spot area). log2(fold change) is calculated with SCR MACs as reference. Differences of means between groups were tested using one-way ANOVA with repeated measures followed by Dunnett’s post-hoc test. N=3/group

MIP1α: ANOVA F(3,6)=5.935, P-value=0.0315, followed by Dunnett’s post-hoc tests: ES40KD-SCR=-0.2016 [-0.3663, -0.0368], P-value40KD-SCR=0.0219; ES41KD-SCR=-0.1603 [-0.3250, 0.0044], P-value_41KD-SCR_=0.0555; ES_DKD-SCR_=-0.0632 [-0.2279, 0.1015], P- value_DKD-SCR_=0.5377.

MIF: ANOVA F(3,6)=6.201, P-value=0.0287, followed by Dunnett’s post-hoc tests: ES40KD-SCR=-0.2372 [-0.4769, 0.00257], P-value40KD-SCR=0.0521; ES41KD-SCR=-0.2244 [-0.4641, 0.0152], P-value_41KD-SCR_=0.0640; ES_DKD-SCR_=-0.3173 [-0.5570, -0.0773], P- value_DKD-SCR_=0.0154.

IL8: ANOVA F(3,6)=2.734, P-value=0.1361, followed by Dunnett’s post-hoc tests: ES40KD- SCR=-0.1798 [-0.3958, 0.0363], P-value40KD-SCR=0.0965; ES41KD-SCR=-0.0678 [-0.2840, 0.1482], P-value41KD-SCR=0.6653; ESDKD-SCR=-0.1504 [-0.3665, 0.0655], P-value_DKD-SCR_=0.1661.

IL1RA: ANOVA F(3,6)=32.74, P-value=0.0004, followed by Dunnett’s post-hoc tests: ES40KD-SCR=-0.5182 [-0.7117, -0.3246], P-value40KD-SCR=0.0004; ES41KD-SCR=-0.5519 [-0.7454, -0.3583], P-value_41KD-SCR_=0.0003; ES_DKD-SCR_=-0.3719 [-0.5655, -0.1783], P- value_DKD-SCR_=0.0025.

IL1β: ANOVA F(3,6)=2.989, P-value=0.1177, followed by Dunnett’s post-hoc tests: ES40KD-SCR=-0.2059 [-0.9746, 0.5628], P-value40KD-SCR=0.7523; ES41KD-SCR=-0.1785 [-0.9472, 0.5902], P-value_41KD-SCR_=0.8159; ES_DKD-SCR_=-0.7064 [-1.475, 0.0623], P- value_DKD-SCR_=0.0684.

ICAM1: ANOVA F(3,6)=14.42 P-value=0.0038, followed by Dunnett’s post-hoc tests: ES40KD-SCR=-0.1981 [-0.2943, -0.1019], P-value40KD-SCR=0.0017; ES41KD-SCR=-0.1339 [-0.2301, -0.0375], P-value_41KD-SCR_=0.0122; ES_DKD-SCR_=-0.1339 [-0.2301, -0.0377], P- value_DKD-SCR_=0.0122.

CXCL11: ANOVA F(3,6)=4.333 P-value=0.0601, followed by Dunnett’s post-hoc tests: ES40KD-SCR=-0.1663 [-0.3380, 0.0054], P-value40KD-SCR=0.0565; ES41KD-SCR=-0.1148 [-0.2865, 0.0568], P-value_41KD-SCR_=0.1856; ES_DKD-SCR_=-0.1790 [-0.3508, -0.0072], P- value_DKD-SCR_=0.0425.

CCL5: ANOVA F(3,6)=15.64 P-value=0.0031, followed by Dunnett’s post-hoc tests: ES40KD-SCR=-0.1876 [-0.2799, -0.0953], P-value40KD-SCR=0.0018; ES41KD-SCR=-0.1615 [-0.2537, -0.0692], P-value_41KD-SCR_=0.0040; ES_DKD-SCR_=-0.1293 [-0.2216, -0.0371], P- value_DKD-SCR_=0.0119.

TNFα: ANOVA F(3,6)=93.00 P-value<0.0001, followed by Dunnett’s post-hoc tests: ES40KD-SCR=-0.6826 [-0.9495, -0.4157], P-value40KD-SCR=0.0005; ES41KD-SCR=-0.7145 [-

0.9814, -0.4476], P-value41KD-SCR=0.0004; ESDKD-SCR=-1.438 [-1.705, -1.171], P-value_DKD-SCR_<0.0001.

SERPINE1: ANOVA F(3,6)=8.744 P-value=0.0131, followed by Dunnett’s post-hoc tests: ES40KD-SCR=-0.2327 [-0.3997, -0.0657], P-value40KD-SCR=0.0122; ES41KD-SCR=-0.0641 [-0.2312, 0.1028], P-value_41KD-SCR_=0.5366; ES_DKD-SCR_=-0.2097 [-0.3766, -0.0427], P- value_DKD-SCR_=0.0196.

Effect sizes (ES) are reported as unstandardized point estimates with 95% confidence intervals in the same unit as depicted on each graph

### Supplementary Table legends

**Supplementary Table 1**

**LAM_datasets sheet.** Human and mouse lipid-associated macrophage (LAM) sc/snRNA-seq datasets used to reconstruct gene regulatory networks.

**LAM_genesets sheet.** Human and mouse LAM genesets used in regulon enrichment, RRHO, and other analyses (genes listed in alphabetical order).

**LAM_TFs sheet.** Candidate human and mouse LAM TFs nominated in more than half human and mouse GRNs, respectively (listed in alphabetical order).

**Supplementary Table 2**

**BHLHE40/41_regulons sheet.** BHLHE40 and BHLHE41 human and mouse regulons from meta-analysed human and mouse GRNs. Meta-analyzed GRNs were generated by aggregating all the bootstraps from individual networks generated by ARACNe.

**LAM_genes_in_BHLHE40/41_regulons sheet.** Human and mouse LAM genes in BHLHE40 and BHLHE41 human and mouse regulons (see BHLHE40/41_regulons sheet). Human LAM genes in BHLHE40 and BHLHE41 human regulons. LAM genes were selected from Jaitin *et al*. [3] (Dataset S6, FDR Adj.P-value < 0.05). Mouse LAM genes in Bhlhe40 and Bhlhe41 mouse regulons. LAM genes were selected from Keren-Shaul *et al*. [10] (Table S3, FDR Adj.P-value < 0.05).

**LAM_genes_with_promoter_proxy-bound_by_BHLHE40/41 sheet.** Human lipid-associated macrophage (LAM) genes with promoters proxy-bound by BHLHE40 and/or BHLHE41. LAM genes were selected from Jaitin *et al*. [3] (Dataset S6, FDR Adj.P-value < 0.05).

**Genes_in_most_significant-RRHO_overlaps sheet.** Genes in most significant overlaps from rank-rank hypergeometric overlap (RRHO) analyses. Z scores (Z.std) from RNA-seq differential gene expression analyses were assigned to each overlapping gene to run Ingenuity Pathway Analysis (IPA). Human LAM signature genes in Figure 4A and Supplementary Figure 7A were selected from Jaitin *et al.* [3] (Dataset S6, FDR Adj.P-value < 0.05). Mouse LAM signature genes in Figure 7A were selected from Keren-Shaul *et al.* [10] (Table S3, FDR Adj.P-value < 0.05).

Supplementary Table 3. Differential gene expression (DGEA) and gene set enrichment (GSEA) analysis of *BHLHE40* and/or *BHLHE41* knockout (KO) in human iPSC-derived microglia (iMGLs). 40KO = BHLHE40 KO iMGLs, 41KO = BHLHE41 KO iMGLs, DKO = BHLHE40 and BHLHE41 double KO iMGLs, WT = iMGLs derived from the parental iPSC line.

Supplementary Table 4. Differential gene expression (DGEA) and gene set enrichment (GSEA) analysis of *BHLHE40* and/or *BHLHE41* knockdown (KD) in human THP-1 macrophages (MACs). 40KD = MACs treated with BHLHE40 siRNA, 41KD = MACs treated with BHLHE41 siRNA, DKD = MACs treated with BHLHE40 and BHLHE41 siRNA, SCR = MACs treated with scrambled siRNA.

Supplementary Table 5. Differential gene expression (DGEA) and gene set enrichment (GSEA) analysis of *Bhlhe40* and *Bhlhe41* double knockout (DKO) in mouse microglia. DKO = Bhlhe40/41 DKO mouse microglia, compared to microglia derived from wild-type control mice.

Supplementary Table 6. Sequences of primers used for RT-qPCR and sequences of guide RNAs, single-stranded oligodeoxynucleotides (ssODNs), and PCR primers used for CRISPR/Cas9-mediated HDR.

